# Ciliated keratinocytes define a spatially restricted pro-inflammatory state in psoriasis and atopic dermatitis

**DOI:** 10.64898/2026.11.11.766039

**Authors:** Mikihito Ike, Yoshiaki Hayashi, Motoki Nakamura, Haoyuan He, Ayane Nakaoku, Ken J. Ishii, Fumitaka Fujita, Daisuke Motooka, Manami Toriyama, Akimichi Morita

**Affiliations:** Department of Pharmaceutical Sciences, The University of Osaka, Suita, Japan; Research Institute for Microbial Diseases (RIMD), The University of Osaka, Suita, Japan; Department of Geriatric and Environmental Dermatology, Nagoya City University Graduate School of Medical Sciences, Nagoya, Japan; Division of Vaccine Science, Department of Microbiology and Immunology, The University of Tokyo, Tokyo, Japan; International Vaccine Design Center, The Institute of Medical Science (IMSUT), The University of Tokyo, Tokyo, Japan; Mandom Corporation, Osaka, Japan

**Author notes:** Corresponding author: Manami Toriyama, Research alliance laboratory for advanced cosmetic science Graduate School of Pharmaceutical Sciences, The University of Osaka, 1-6 Yamadaoka, Suita, Osaka, Japan 565-0871 Suita, Osaka, Japan, Akimichi Morita, Department of Geriatric and Environmental Dermatology Nagoya City University Graduate School of Medical Sciences, 1 Kawasumi, Mizuho-cho, Mizuho-ku, Nagoya, Aichi, Japan 467-8602 Nagoya, Aichi, Japan.

## Abstract

**Background:** Psoriasis vulgaris (Pso) and atopic dermatitis (AD) are characterized by recurrent flares and incomplete disease resolution. Primary cilia are antenna-like sensory organelles that project from the cell surface and coordinate extracellular signaling. Although epidermal ciliated cells are rarely detected in healthy skin and are selectively increased in AD and Pso, their molecular identity, functional properties, and clinical relevance remain unclear.

**Objective:** To characterize epidermal ciliated cells in Pso and AD and determine their molecular identity, functional properties, and clinical relevance.

**Methods:** We integrated single-cell-resolved spatial transcriptomics with primary cilia immunostaining in healthy skin (n=6), AD lesions (n=10), and paired psoriatic skin before and after phototherapy (n=7 each). Functional analyses using HaCaT cells with increased primary cilia formation included MAPK/NF-κB signaling assays, ATAC-seq, and pathway inhibition assays.

**Results:** Ciliated cells were largely restricted to AD and both pre- and post-treatment psoriatic skin, with KCs accounting for more than 70% of all ciliated cells. Ciliated KCs were predominantly localized to the basal and spinous layers. These cells exhibited disease-and stage–specific inflammatory programs and formed spatially organized communication networks with surrounding cells, including non-ciliated KCs, fibroblasts, endothelial cells, and myeloid cells, suggesting a ciliated KC-centered inflammation niche. Mechanistically, increased primary cilia formation enhanced MAPK and NFκB signaling and increased chromatin accessibility at inflammatory gene loci enriched for AP-1 motifs, even in the absence of stimulation, consistent with a chromatin-primed state. Higher residual ciliated KC abundance after phototherapy was associated with subsequent treatment escalation. ERK1/2 inhibition reduced cytokine-induced primary cilia formation.

**Conclusions:** Ciliated KCs represent an inflammatory epithelial cell state associated with residual disease activity in AD and Pso. The suppression of cytokine-induced primary cilia formation by ERK1/2 inhibition suggests that inflammation-associated–primary cilia formation may be modifiable process in persistent skin inflammation.

**Lay summary:** Psoriasis (Pso) and atopic dermatitis (AD) are common chronic inflammatory skin diseases. They can cause redness, itching, scaling, and repeated flare-ups. Treatments such as steroids, biologics, and phototherapy can improve skin symptoms, but inflammation-related changes may remain in the skin even after the skin appears clinically improved.

In this study, we focused on skin cells that carry antenna-like structures called primary cilia. Primary cilia help cells sense signals from their surroundings. Although these structures are rarely seen in healthy skin, we found that ciliated cells were increased in the skin of patients with AD and Pso, most of them were keratinocytes (KCs), the main cells of the outer layer of the skin. Using spatial transcriptomics analysis combined with immunostaining, we examined where these cells were located and their gene expression patterns. Ciliated KCs showed inflammatory signaling, communication with surrounding cells, and changes in chromatin, the structure that controls how genes are switched on and off. These changes were linked to AP-1, a group of factors involved in inflammation-related gene activity. We also found that ciliated KCs remained in psoriasis skin samples after phototherapy and may be associated with later treatment escalation. Blocking ERK1/2 signaling reduced inflammation-induced primary cilia formation.

These findings suggest that ciliated KCs may contribute to persistent skin inflammation. They also suggest that targeting inflammation-associated primary cilia formation could be explored as a future approach to limit persistent skin inflammation.

*What is already known about this topic?:*

- Atopic dermatitis and psoriasis are chronic, relapsing inflammatory skin diseases. Although corticosteroids, biologics, and phototherapy can improve clinical symptoms, disease persistence and recurrence remain major clinical challenges.
- Primary cilia are signaling organelles that sense extracellular cues and coordinate cellular responses.
- Epidermal ciliated cells are rare in healthy skin but are increased in the epidermis of patients with atopic dermatitis and psoriasis.

*What does this study add?:*

- Epidermal ciliated cells in atopic dermatitis and psoriasis were identified mainly as keratinocytes (KCs), a disease-associated epithelial population rare in healthy skin.
- Ciliated KCs showed disease- and treatment-stage-specific inflammatory programs, spatial communication with surrounding epidermal, stromal, vascular, and immune cells, and persistence after phototherapy.
- Enhanced primary cilia formation was linked to MAPK/NFkB signaling and AP-1-enriched inflammatory chromatin accessibility, and ERK1/2 signaling regulated cytokine-induced primary cilia formation.

*What is the translational message?:*

- Residual ciliated keratinocytes (KCs) may represent persistent molecular disease activity that is not fully captured by clinical severity scores.
- Detection of ciliated KCs may help identify patients at risk of disease persistence, recurrence, or subsequent treatment escalation.
- ERK1/2 inhibition suppressed inflammation-induced primary cilia formation, suggesting that inflammation-associated ciliogenesis may be a modifiable process in atopic dermatitis and psoriasis.

## INTRODUCTION

Atopic dermatitis (AD) and Psoriasis vulgaris (Pso) are common chronic relapsing inflammatory skin diseases.^1–3^ Despite advances in steroids, biologics, and phototherapy, inflammatory gene expression and transcriptional programs are not fully normalized within the lesional microenvironment after treatment.^4–6^ The molecular basis for this discrepancy between clinical remission and persistent molecular activity remains poorly understood.

Keratinocytes (KCs), the predominant epidermal cell type, are critical regulators of cutaneous inflammation. Although Pso and AD have traditionally been viewed as immune cell-driven diseases, growing evidence indicates that KCs actively shape the local inflammatory microenvironment through cytokine and chemokine production and bidirectional interaction with immune cells, fibroblasts (FBs), and endothelial cells (ECs).^7,8^ However, how KCs integrate diverse signaling inputs to sustain chronic inflammation remains unclear.

Primary cilia are microtubule-based organelles that function as signaling hubs for multiple receptor pathways.^9,10^ They are rarely detected in healthy epidermis but are significantly increased in the epidermis of AD and Pso, and can be induced by Th2- and Th17-cytokines in KCs.^11^ However, the molecular features and inflammatory roles of ciliated KCs remain poorly understood, partly because these rare cells are difficult to isolate.^12^ Here, we combined spatial transcriptomics with primary cilia immunostaining at single-cell resolution in healthy skin, AD, and Pso before (Pso-pre) and after (Pso-post) treatment. We further established KC cell lines that constitutively form primary cilia and performed ATAC-seq to assess chromatin accessibility at inflammatory gene loci. Our findings suggest a mechanism for disease persistence in chronic inflammatory skin diseases and support a framework in which cellular organelles regulate immune responsiveness through epigenomic remodeling.

## MATERIALS AND METHODS

### Study approval

The collection of samples from patients with AD, Pso, HC, and the use of formalin-fixed, paraffin-embedded blocks from prior diagnostic biopsy samples was approved by the Human Research Ethics Committees of the Nagoya City University Clinical Trial Management Center (nos. 60-18-0003, 70-23-0014) and the University of Osaka (nos. yakuhito2019-01, 2019-01-2, 2024-4), and all participants provided written informed consent to participate in this study.

### ST data tissue preparation, generation, preprocessing, and analysis

ST analysis was performed using the 10x Genomics Visium HD Spatial Gene Expression platform on human adult skin tissue samples (Supplementary Table S1). Primary cilia were detected by immunofluorescence staining for acetylated tubulin and pericentrin on serial tissue sections and integrated with the ST section to identify ciliated cells. Cell segmentation was performed using StarDist on the Hoechst-stained nuclear channel, and transcript counts from 2-μm bins were aggregated into single-cell profiles.

### Bulk ATAC-seq cell suspension, data generation, processing, and analysis

Bulk ATAC-seq was performed on mock, ARL13B-overexpressing, and GPR161-overexpressing HaCaT cells using an ATAC-seq kit according to the manufacturer’s instructions. Paired-end reads were trimmed and aligned to the human hg38 reference genome using Bowtie2, and duplicate reads were removed using SAMtools. Further materials and methods are detailed in Appendix S1 (see Supporting Information).

## RESULTS

### Spatial transcriptomics analysis revealed disease-specific gene expression in ciliated KCs

To characterize disease-associated ciliated cells and define their identities in chronic inflammatory skin diseases, we integrated spatial transcriptomics (ST) with immunofluorescence imaging (Figure 1a). Formalin-fixed paraffin-embedded skin samples were obtained from healthy controls (HCs, n = 5), patients with AD (AD, n = 10), and patients with Pso (pso-pre, n = 7), with paired post-phototherapy samples (pso-post, n = 7). Phototherapy efficacy was evaluated by epidermal thickness and Psoriasis Area and Severity Index (PASI) scores (Supplementary Figure S1a; see Supporting information). Serial sections (6, 3, and 6 µm) were immunostained for primary cilia, and the central 3-µm section was also subjected to single-cell-resolved ST (Figure 1a). Three-dimensional integration of serial sections enabled accurate detection of primary cilia in ST samples (Figure 1b and Figure S1b and c; see Supporting information)

**Figure 1.**
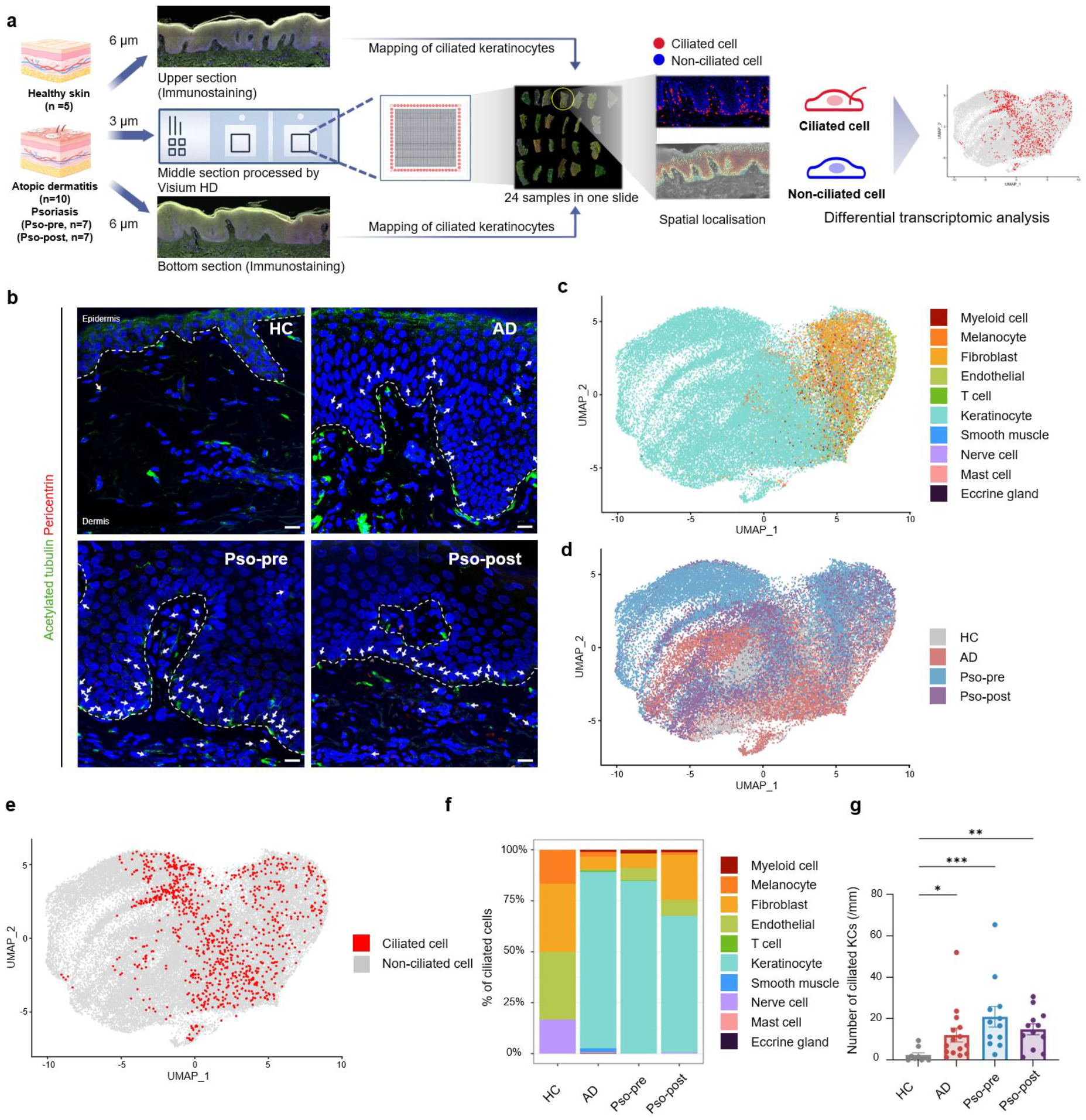
Spatial transcriptomics identifies disease-specific features of ciliated cells. (**a**) Schematic overview of the experimental design and spatial transcriptomic workflow used to identify primary cilium-positive cells (ciliated cells) in human skin by immunofluorescence and to characterize their transcriptional profiles. Illustrations were generated using BioRender (https://biorender.com). (**b**) Immunofluorescence images of primary cilia in human skin. Ciliary axonemes were labeled with acetylated α-tubulin (green), centrosomes with pericentrin (red), and nuclei were stained with Hoechst 33342 (blue). White arrows indicate primary cilia. Scale bar: 15 µm. (**c**) UMAP visualization of 46,033 cells derived from healthy skin (HC), atopic dermatitis (AD), psoriasis before treatment (Pso-pre), and psoriasis after treatment (Pso-post) colored according to annotated cell types. (**d**) UMAP embedding shown in (**c**), colored by disease condition. (**e**) The projection of ciliated cells onto the UMAP embedding is shown, highlighting their distribution across annotated cell populations. (**f**) The bar plot shows the proportion of ciliated cells among the annotated cell types for each disease condition. (**g**) Quantification of epidermal ciliated cells per millimeter of epidermal length across disease conditions. Statistical significance was assessed using one-way ANOVA, followed by Dunnett’s multiple comparison test for comparison with the control group. \**P* < 0.05, \*\**P* < 0.01, \*\*\**P* < 0.001.

We analyzed 29 skin sections and obtained 46,033 cells after quality filtering. Dimensionality reduction and clustering identified 10 distinct cell types, validated by canonical markers and spatial localization (Figure 1c and Figure S1d and e; see Supporting information).^13^ KCs from the AD and Pso formed disease-specific clusters distinct from healthy KCs, and most Pso-post KCs remained closer to AD and Pso-pre KCs than to healthy KCs (Figure 1d).

Ciliated cells were rare in healthy skin but increased in diseased skin across several cell types. Notably, epidermal KCs accounted for more than 70% of all ciliated cells (Figure 1c–f), consistent with the significant increase in epidermal cilia in AD, Pso-pre, and Pso-post compared with HC (Figure 1g). Importantly, ciliated KCs persisted even after clinical improvement (Figure S1a; see Supporting information), suggesting functional roles beyond active disease states. To further understand their functional properties, we performed a detailed analysis of ciliated KCs.

### Ciliated KCs define distinct and disease-specific transcriptional states

We next reanalyzed the ciliated KC subset. UMAP visualization showed that diseased KCs were distinct from healthy KCs (Figure 2a). Because KC maturation differs across epidermal layer, we annotated basal, spinous, and granular populations using canonical markers (Figure 2b and Figure S2a; see Supporting information).^14^ These annotations matched epidermal localization.^13^ Consistent with immunofluorescence imaging, most ciliated KCs localized to basal and spinous clusters (Figure 2c). Ciliated KCs were mainly distributed in the basal layer in the Pso-pre, Pso-post, and AD groups (72%, 65%, and 49%, respectively) and in the spinous layer (26%, 35%, and 51%, respectively), with few in the granular layer (Figure 2a–d).

**Figure 2.**
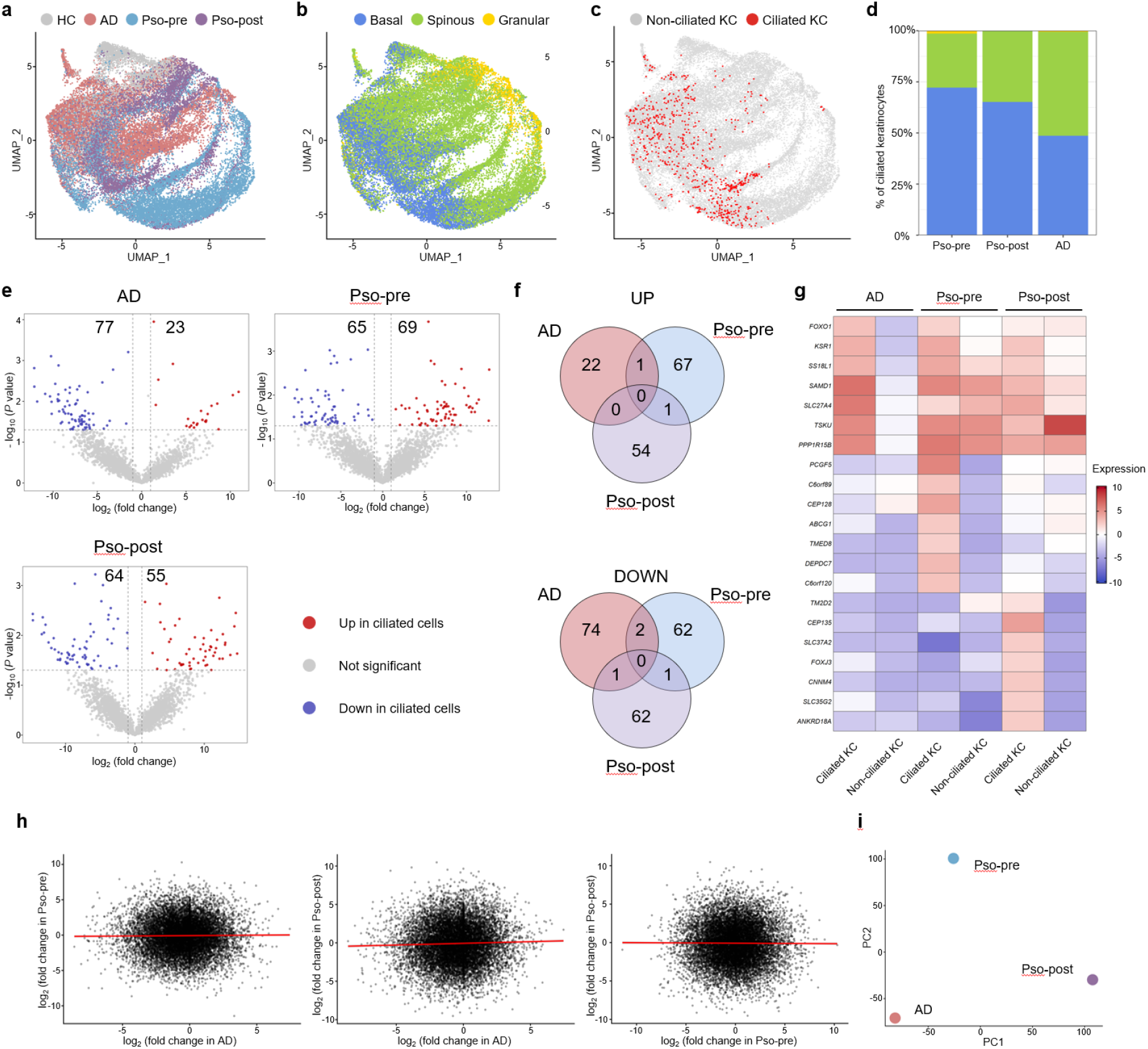
Disease-specific heterogeneity of ciliated KCs in basal and spinous layers. (**a**) UMAP of re-clustered KCs extracted from Fig. 1C, colored according to the disease conditions. (**b**) UMAP embedding is shown in (**a**) annotated with predicted epidermal layer identities based on previously reported layer-specific markers in healthy skin.^12^ (**c**) Projection of ciliated KCs onto the UMAP embedding shown in (**a**). (**d**) Proportion of ciliated KCs in each KC subcluster corresponding to distinct epidermal layers. (**e**)Volcano plots of pseudobulk differentially expressed genes (DEGs) showing upregulated or downregulated genes in ciliated KCs relative to non-ciliated KCs in lesional AD, Pso-pre, and Pso-post. Patients were treated as biological replicates, and DEGs were identified using a paired design, controlling for patient identity. The DEG lists are shown (Supplementary Table S2). (**f**) Venn diagrams showing the overlap of upregulated (top) and downregulated (bottom) genes in ciliated KCs relative to non-ciliated KCs across disease conditions. (**g**) Pseudo-bulk heatmap showing the scaled expression of the top seven DEGs identified in ciliated versus non-ciliated KCs for each disease condition. (**h**) Gene-wise effect sizes (log2 fold change, ciliated vs. non-ciliated) were compared across the disease conditions. Left, AD vs. pso-pre; middle, AD vs. pso-post; right, pso-pre vs. pso-post. Pearson (r) and Spearman (ρ) correlation coefficients are indicated (AD vs Pso pre: r = 0.0096, *P* = 2.06 × 10⁻¹; ρ = 0.0170; AD vs Pso-post: r = 0.0314, *P* = 5.26 × 10⁻⁵; ρ = 0.0270; Pso pre vs Pso post: r = −0.0060, *P* = 4.31 × 10⁻¹; ρ = −0.0098). (**i**) Principal component analysis (PCA) of gene-wise effect sizes (log2 fold change, ciliated vs. non-ciliated) across disease conditions. Each point represents a disease group (AD, Pso-pre, or Pso-post), with PC1 and PC2 capturing the major sources of variation.

To define primary cilia-associated transcriptional changes within each disease context, we compared ciliated KCs with layer-matched non-ciliated KCs in AD, Pso-pre, and Pso-post samples using pseudobulk DEG analysis (Figure S2b and Table S2; see Supporting information). Ciliated KCs showed distinct up- and downregulated gene sets relative to non-ciliated KCs in each condition (Figure 2e), with minimal overlap among AD, Pso-pre, and Pso-post groups (Figure 2f). These results indicate that ciliated KCs acquire disease-specific transcriptional programs rather than a uniform ciliation-associated signature.

Heat map analysis showed that AD upregulated genes were also increased in Pso-pre and Pso-post, although at lower levels, whereas Pso-pre and Pso-post exhibited largely distinct profiles (Figure 2g). Thus, AD ciliated KCs may represent a shared activation state, whereas Pso ciliated KCs acquire additional disease-specific programs. *FOXO1* and *KSR1*, associated with MAPK signaling, were upregulated across conditions,^15, 16^ whereas centrosome-associated *CEP128* and *CEP135* were selectively enriched in Pso, suggesting enhanced ciliogenesis-related programs (Figure 2g).^17,18^

Gene-wise effect sizes showed no significant correlations between disease conditions (Figure 2h), even among significant DEGs (Figure S2c; see Supporting information), and principal component analysis (PCA) clearly separated AD, Pso-pre, and Pso-post (Figure 2i). Together, these results indicate that ciliated KCs form disease-specific rather than conserved transcriptional states.

### Ciliated KCs exhibit environment-dependent functional states in Pso

Given the distinct transcriptional profiles of ciliated KCs in Pso-pre and Pso-post samples, we next analyzed paired samples to define treatment-associated changes in their functional states. Gene Ontology (GO) analysis confirmed enrichment of “non-motile cilium assembly” in both conditions, supporting their classification (Figure 3a and b and Table S3; see Supporting information). Pso-pre ciliated KCs were enriched for energy metabolism and intracellular transport pathways, whereas Pso-post ciliated KCs showed angiogenesis-related enrichment (Figure 3a and b), suggesting that ciliated KCs adopt distinct functional programs in active versus post-treatment psoriatic skin.

**Figure 3.**
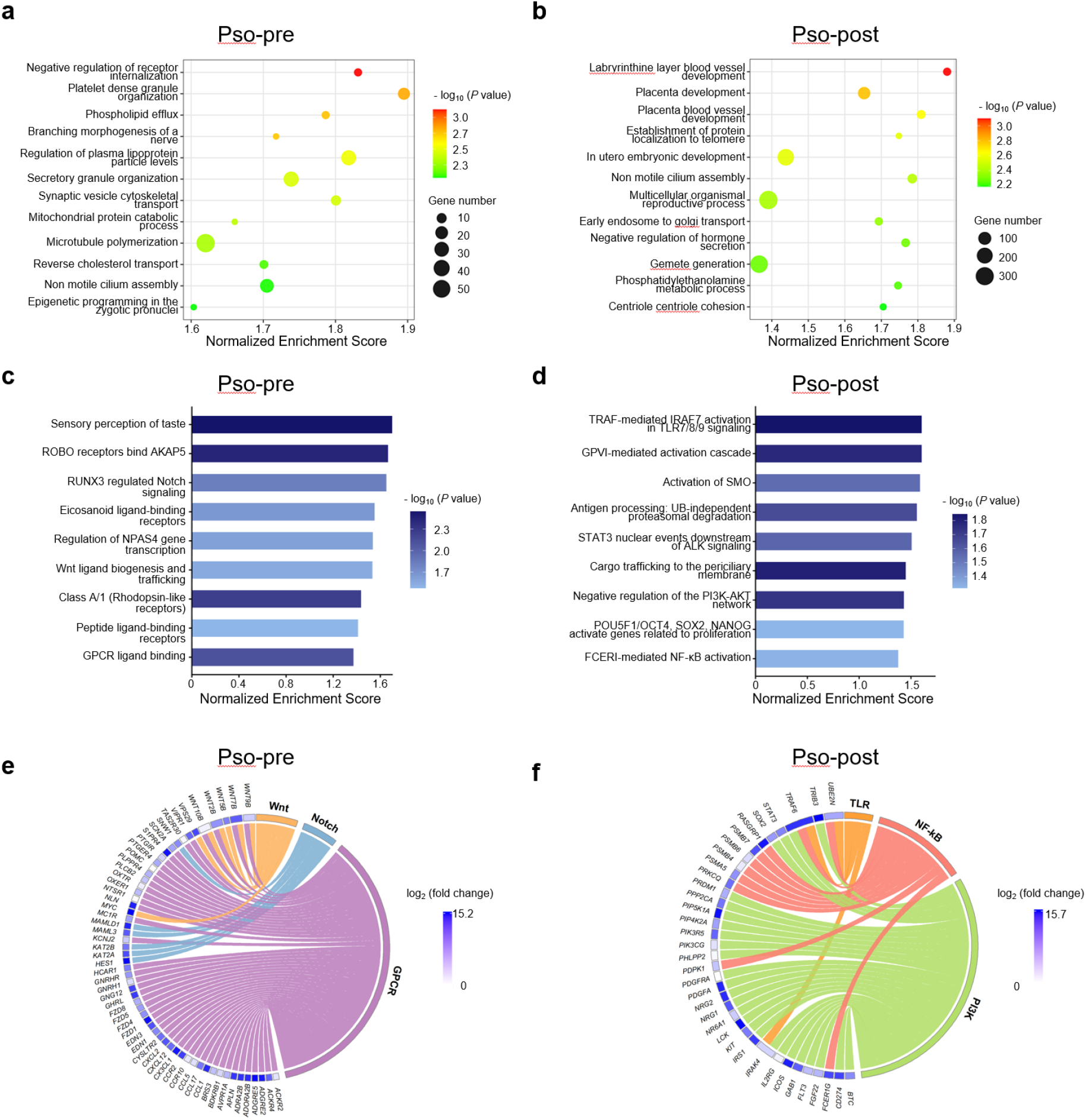
Signaling pathways enriched in ciliated KCs in Pso-pre and Pso-post tissues. (**a**) Gene Ontology (GO) analysis of genes upregulated in ciliated KCs relative to non-ciliated KCs in Pso-pre-samples. (**b**) GO analysis of genes upregulated in ciliated KCs relative to non-ciliated KCs in pso-post-samples. (**c**) Bar plot showing significantly enriched signaling pathways identified by Reactome analysis of genes upregulated in ciliated KCs relative to non-ciliated KCs in Pso-pre samples. (**d**) Bar plot showing significantly enriched signaling pathways identified by Reactome analysis of genes upregulated in ciliated KCs relative to non-ciliated KCs in Pso-post samples. (**e**) Circos plot illustrating selected signaling pathways (Wnt, Notch, and GPCR) enriched in ciliated KCs relative to non-ciliated KCs in Pso-pre samples, based on GSEA results shown in (**c**) and their associated DEGs. (**f**) Circos plot illustrating selected signaling pathways (TLR, NFκB, and PI3K) enriched in ciliated KCs relative to non-ciliated KCs in Pso-post samples, based on GSEA results shown in (**d**), and their associated DEGs.

Reactome analysis also revealed distinct signaling programs (Table S4; see Supporting information). In Pso-pre, ciliated KCs were enriched for GPCR, Wnt, and Notch signaling (Figure 3c), with increased expression of Notch- and Wnt-related genes involved in KC proliferation and differentiation (Figure 3e).^19–22^ Chemokines were also upregulated, suggesting roles for ciliated KCs in immune cell recruitment and the formation of an inflammatory microenvironment.^23–25^

In Pso-post, ciliated KCs were enriched for TLR– NF-κB, STAT3, and PI3K–AKT signaling (Figure 3d). Upregulation of innate immune genes (*TRAF6*, *IRAK4*, *UBE2N*) and survival-related genes (*PIK3CG*, *PDPK1*, *PDGFRA*, *IRS1*, and *STAT3*) (Figure 3f) suggested persistent inflammatory and survival signaling even after treatment.^26–29^

Thus, ciliated KCs adopt environment-dependent functional state. In Pso-pre, they are associated with proliferation–differentiation imbalance,^21,22^ and to inflammatory microenvironment formation, whereas in Pso-post, they maintain innate immune and survival programs that may support inflammatory priming.

### Spatially resolved cell–cell communication networks involve ciliated KCs in Pso

To examine how ciliated KCs communicate with surrounding cells, we defined cells within 50 µm as adjacent and those within 50–100 µm as distal (Figure 4a). Across both distance ranges and disease conditions, KCs, FBs, and ECs together accounted for more than half of the local cell population (Figure 4b). Next, we analyzed ligand receptor interactions centered on ciliated KCs using CellPhoneDB (Table S5; see Supporting information). In the Pso-pre samples, ciliated KCs showed the highest number of interactions with both ciliated and non-ciliated KCs, followed by FBs, ECs, and myeloid cells (Figure 4c). Notably, similar interaction patterns were observed in the Pso-post samples, although interactions with myeloid cells were more prominent than in Pso-pre samples. This finding suggested a possible link between ciliated KCs and vascular-associated inflammatory niches. Since Pso severity has been reported to correlate with the number of dermal papillary microvessels,^30^ we examined the spatial relationship between ciliated KCs and ECs. Ciliated KCs were located significantly closer to CD31-positive cells than non-ciliated basal KCs (Figure 4d and e), supporting a spatial basis for ciliated KCs –ECs communication.

**Figure 4.**
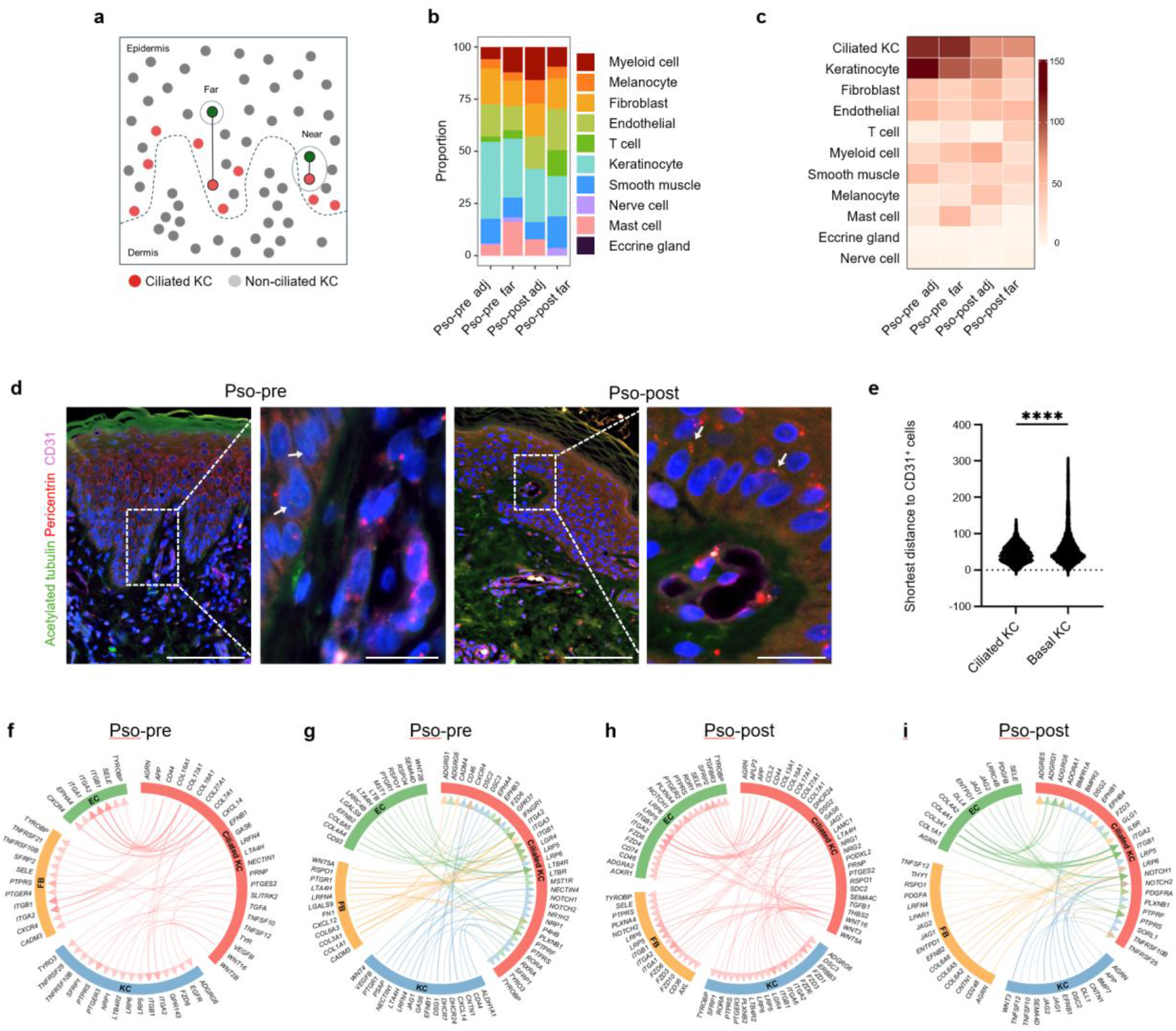
Spatially resolved ligand-receptor interactions of ciliated KCs in Pso. (**a**) Schematic illustrating the definition of adjacent (‘near,’ ≤50 μm) and distal (‘far,’ 50–100 μm) cells relative to ciliated KCs. (**b**) Proportion of cell types interacting with ciliated KCs based on ligand–receptor interaction analysis. (**c**) Heatmaps showing the number of adjacent ligand–receptor pairs in Pso-pre samples. (**d**)Immunofluorescence images showing the spatial relationship between CD31-positive ECs (CD31, magenta) and ciliated cells in Pso-pre and Pso-post skin. Primary cilia were labeled with acetylated α-tubulin (green) and basal bodies were labeled with pericentrin (red). Scale bars, 20 µm (left), 100 µm (middle left). (**e**) Violin plot showing the shortest distance between ciliated KCs and CD31-positive cells compared to non-ciliated basal KCs. Comparisons between the two groups were performed using an unpaired two-tailed Student’s t-test. \*\*\*\**P* < 0.0001. (**f**) Circos plot illustrating adjacent ligand-receptor interactions originating from ciliated KCs in Pso-pre samples. Ligands specifically expressed by ciliated KCs were predicted to interact with receptors on non-ciliated KCs, FBs, and ECs. (**g**) Circos plot illustrating adjacent ligand-receptor interactions received by ciliated KCs in Pso-pre samples. Ligands derived from non-ciliated KCs, FBs, and ECs were predicted to interact with receptors preferentially expressed in ciliated KCs. (**h**) Circos plot illustrating adjacent ligand–receptor interactions originating from ciliated KCs in Pso-post samples. Ligands specifically expressed by ciliated KCs were predicted to interact with receptors on non-ciliated KCs, FBs, and ECs. (**i**) Circos plot illustrating adjacent ligand-receptor interactions received by ciliated KCs in Pso-post samples. Ligands derived from non-ciliated KCs, FBs, and ECs were predicted to interact with receptors preferentially expressed in ciliated KCs.

We then extracted the major ligand–receptor pairs between ciliated KCs and adjacent non-ciliated KCs, FBs, and ECs (Figure 4f–i). In Pso-pre, ciliated KCs were predicted to initiate communication programs, including *CXCL14*– *CXCR4* with ECs and FBs, *TNFSF10*–*TNFRSF10B* with non-ciliated KCs and FBs, and *TGFA*– *ADGRG6* with non-ciliated KCs (Figure 4f), which were largely restricted to ciliated KCs, suggesting a role for ciliated KCs in tissue remodeling and inflammatory microenvironment formation in psoriatic lesions.^31^ Ciliated KCs were also predicted to preferentially receive *CXCL14* from non-ciliated KCs via *CXCR4* (Figure 4g), indicating that CXCL14–CXCR4 signaling is selectively engaged in ciliated KCs but not in non-ciliated KCs, and may regulate their migratory behavior.^32,33^

In Pso-post, ciliated KCs were predicted to produce *TGFB1*, and *CCL2*, engaging *TGFBR3* and *ACKR1* on ECs (Figure 4h), consistent with roles in vascular regulation and leukocyte recruitment.^34,35^ Furthermore, ciliated KCs retained responsiveness to pro-inflammatory signals, including *TNFSF12*– *TNFRSF25* from non-ciliated KCs and FBs and *TNFSF10*–*TNFRSF10B* from non-ciliated KCs (Figure 4i), indicating persistent signaling activity even after treatment.

Under both pre- and post-treatment conditions, the ciliated KCs exhibited extensive interactions with multiple cell types (Figure 4f–i). Ephrin B–Eph and Collagen–integrin interactions suggested contact-dependent signaling and ECM engagement between ciliated KCs and neighboring cells.^36,37^ Furthermore, Wnt ligands production including *WNT5A*, *WNT2B*, *WNT3*, and *WNT16*, and Notch signaling from adjacent KCs further indicated roles in promoting KC hyperproliferation and differentiation, respectively.^38–40^ Notably, ligand–receptor interactions between ciliated KCs and distant cell populations were largely comparable to those observed between adjacent cells. However, Notch interactions were restricted to adjacent cells (Figure S3; see Supporting information), highlighting spatially confined signaling. Together, these findings suggest that ciliated KCs are positioned at the center of the disrupted balance between proliferation and differentiation characteristics of psoriatic epidermis.

### Ciliated KCs in AD exhibit MAPK–AP-1 activation and interact with neurons

We next compared AD ciliated KCs with layer-matched non-ciliated KCs. GO analysis revealed enrichment of lipid metabolism and environmental response pathways, including fatty acid metabolism and response to chemical stimuli (Figure S4a; see Supporting information), as well as organelle homeostasis and stress responses. Reactome analysis further identified significant enrichment of “Activation of the AP-1 family of transcription factors” (Figure S4b; see Supporting information), and Circos visualization highlighted MAPK–AP-1 related genes, including *MAPK1*, *MAPK8*, *MAPK10*, *MAPK11*, and *JUN* (Figure S4c; see Supporting information). These results suggest enhanced MAPK– AP-1 signaling in AD ciliated KCs.^41^

Cell–cell interaction analysis using CellPhoneDB showed that ciliated KCs interact extensively with KCs, FBs, and ECs, similar to Pso (Figure S4d–f; see Supporting information), and prominently with neuronal cells (Figure S4d; see Supporting information). Ligand–receptor analysis revealed inflammatory interactions from ciliated to non-ciliated KCs, including *JAG1*–*NOTCH3* and *LTA4H*–*LTB4R2*, consistent with those observed in Pso (Figure S4e; see Supporting information). Ciliated KCs both sent and received *LTA4H*–*LTB4R* signals with ECs, non-ciliated KCs, and mast cells, suggesting a role in amplifying inflammatory microenvironments.^42^

WNT16–SFRP1, CLSTN1–NRXN1, and GAS6–AXL interactions further suggested roles for ciliated KCs in neuronal communication and sensory regulation (Figure S4f; see Supporting information).^43–45^ Additional melanocyte-derived *EFNB1*–*EPHB3*/*EPHB6* interactions further supported their involvement in multicellular signaling networks. Together, these results indicate that ciliated KCs in AD exhibit lipid metabolism remodeling, MAPK–AP-1 activation, and broad interactions with non-ciliated KCs, neuronal cells, immune cells, and melanocytes.

### Primary cilia formation enhances MAPK and NF-κB inflammatory signaling in KCs

Reactome and CellPhoneDB analysis suggested the activation of MAPK and NF-κB signaling in ciliated KCs. To further validate this, we generated HaCaT cells, an immortalized human KC cell line, with constitutive cilia formation by overexpressing ARL13B and GPR161 (Figure 5a). ARL13B is essential for primary cilia formation,^46^ whereas GPR161 is a cilia-localized GPCR that activates cAMP– PKA signaling.^47^ Immunofluorescence staining confirmed a significant increase in ciliated cells in ARL13B- and GPR161-overexpressing cells compared with mock controls (Figure 5a and b).

**Figure 5.**
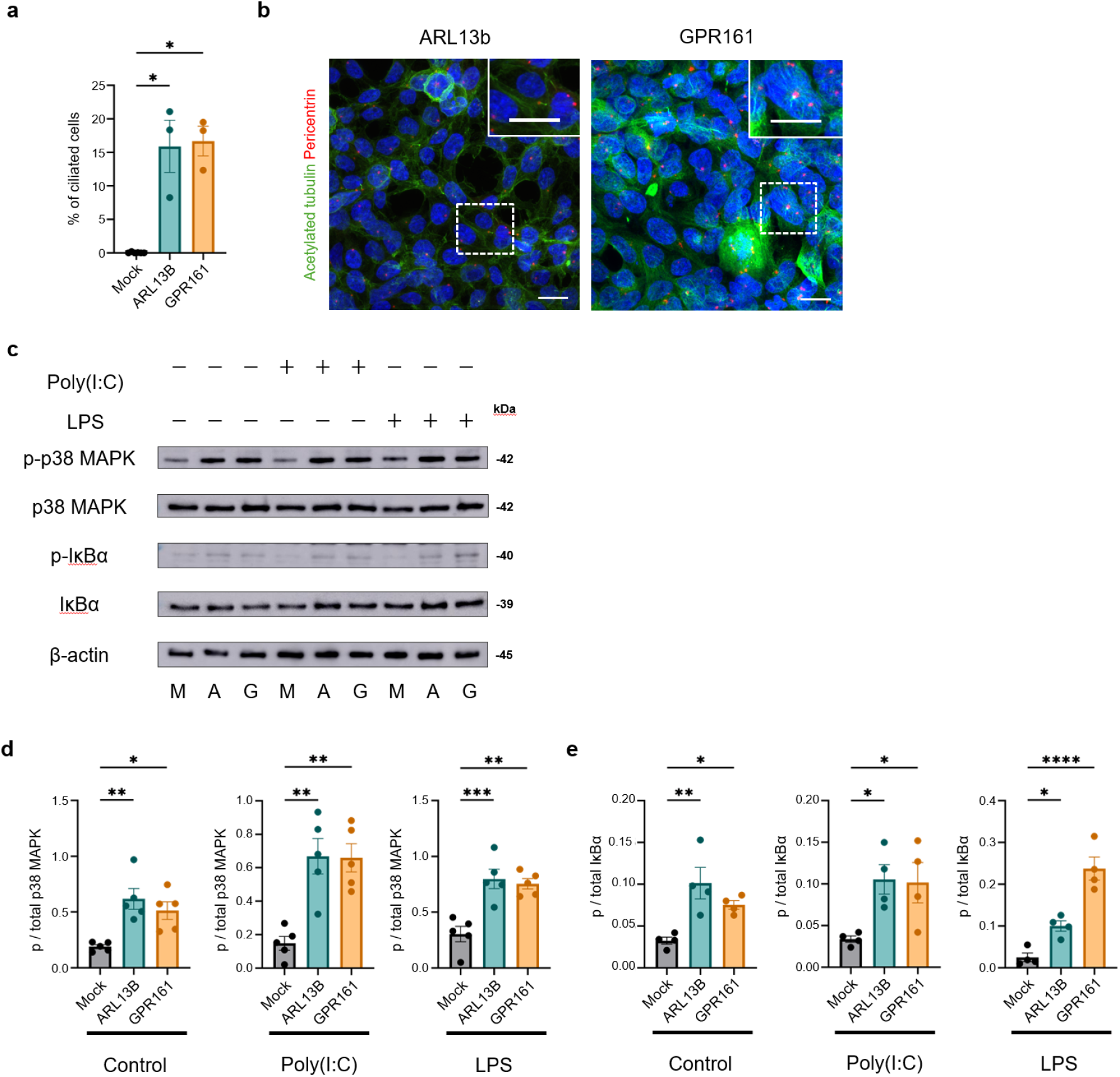
Primary cilia formation promotes p38 MAPK and IκBα activation in HaCaT cells. (**a**) Percentage of ciliated HaCaT cells following overexpression of ARL13B or GPR161 compared to mock-transfected controls. Statistical significance was assessed using one-way ANOVA followed by Dunn’s multiple comparison test versus the control group. \**P* < 0.05. (**b**) Immunofluorescence staining of primary cilia in mock-, ARL13B-, and GPR161-overexpressing HaCaT cells. Primary cilia were labeled with acetylated α-tubulin (green), basal bodies with pericentrin (red), and nuclei with Hoechst 33342 (blue). Scale bar, 15 μm. (**c**) Immunoblotting of phosphorylated and total p38 MAPK and IκBα in mock (M)-, ARL13B (A)-, and GPR161(G)-overexpressing HaCaT cells. The cells were stimulated with 30 μg/mL Poly(I:C) or 10 μg/mL LPS for 15 min. (**d**) Quantification of p38 MAPK activation is shown in (**c**) and is expressed as the ratio of phosphorylated p38 MAPK to total p38 MAPK (p-p38/total p38). Statistical significance was assessed using one-way ANOVA, followed by Dunnett’s multiple comparison test versus the control group. \**P* < 0.05, \*\**P* < 0.01, \*\*\**P* < 0.001. (**e**) Quantification of IκBα phosphorylation is shown in (**c**), expressed as the ratio of phosphorylated IκBα to total IκBα (p-IκBα/total IκBα). Statistical significance was assessed using one-way ANOVA, followed by Dunnett’s multiple comparison test versus the control group. \**P* < 0.05, \*\**P* < 0.01, \*\*\**P* < 0.001, \*\*\*\**P* < 0.0001.

To evaluate the impact of primary cilia on inflammatory signaling, cells were stimulated with poly(I:C) or LPS, and MAPK and NF-κB activation were analyzed. Both ARL13B- and GPR161-overexpressing cells showed significantly increased phosphorylation of p38 MAPK and IκBα compared to mock controls, even in the absence of stimulation (Figure 5c-e). Upon stimulation, mock and ARL13B-overexpressing cells showed no significant changes in p/total IκBα or p/total p38 MAPK ratios.

In contrast, GPR161-overexpressing cells showed enhanced responsiveness, with a significant increase in p/total IκBα following LPS stimulation and a trend toward increased p/total p38 MAPK levels (Figure S5a and b; see Supporting information). These findings suggest that enhanced primary cilia formation elevates basal MAPK and NF-κB activation and may increase responsiveness to inflammatory stimuli.

### Primary cilia formation enhances chromatin accessibility at AP-1 motif-containing inflammatory genes

Enhanced primary cilia formation elevated basal MAPK and NF-κB activation and increased responsiveness to inflammatory stimuli. We therefore asked whether enhanced primary cilia formation is also associated with chromatin-level inflammatory priming and performed bulk ATAC-seq in unstimulated ARL13B- and GPR161-overexpressing HaCaT cells and mock controls. (Figure 6a and Figure S6a and Table S6; see Supporting information). ARL13B overexpression induced differential accessibility at 2,436 regions, including 877 promoters associated with inflammatory genes (Figure 6b). GPR161 overexpression induced a markedly broader chromatin remodeling, with 49,785 differentially accessible regions (DARs) showing increased accessibility, including 4,330 promoters of key inflammatory genes (Figure 6c).

**Figure 6.**
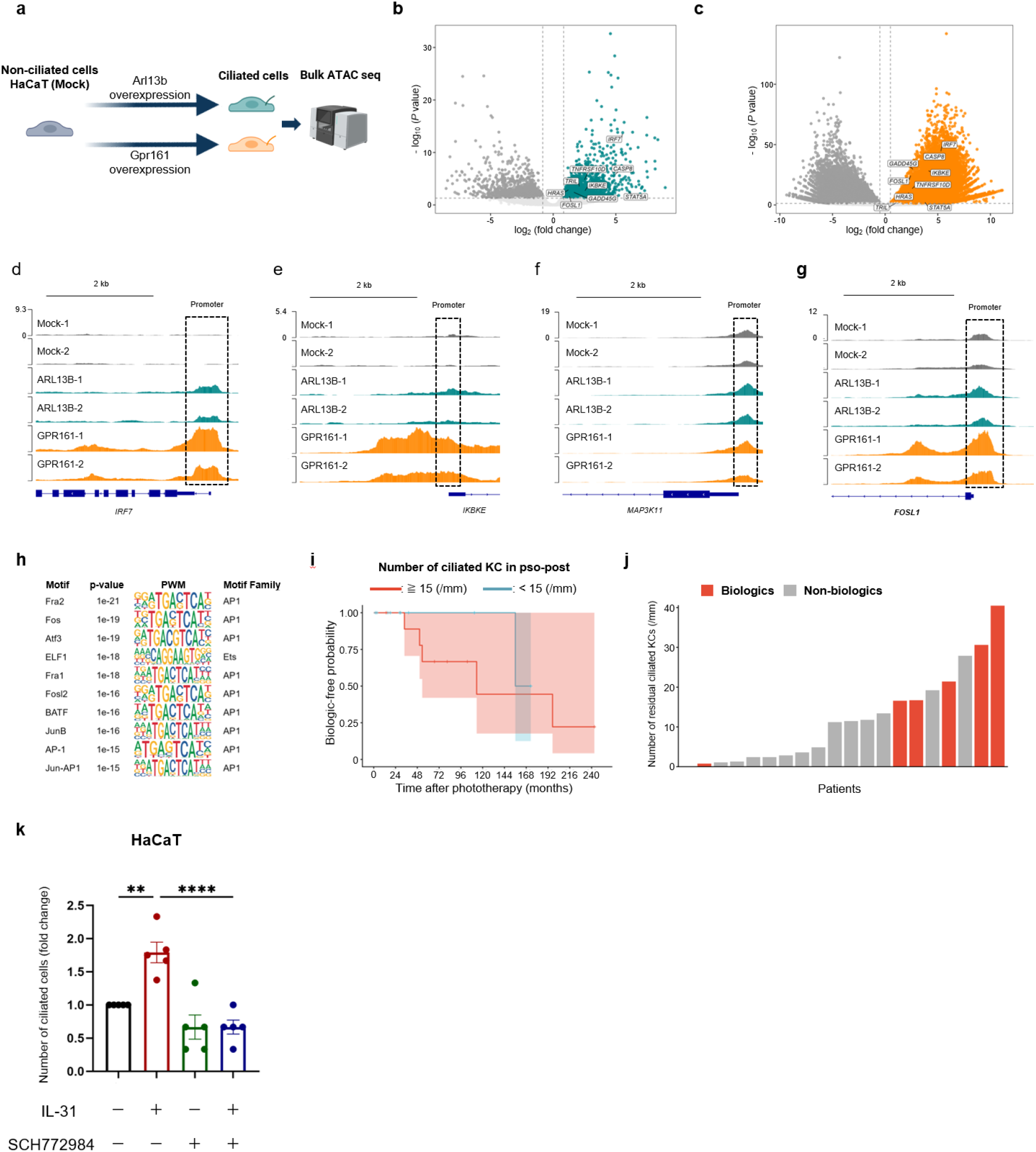
Primary cilia formation induces chromatin remodeling and activates inflammatory gene programs in KCs. (**a**) Schematic overview of the bulk ATAC-seq workflow in HaCaT cells overexpressing ARL13B or GPR161. Illustrations were generated using BioRender (https://biorender.com). (**b**) Volcano plot showing differentially accessible regions (DARs) identified by ATAC-seq in ARL13B-overexpressing HaCaT cells compared to mock controls (n = 3,696 DARs). (**c**) Volcano plots showing DARs identified by ATAC-seq in GPR161-overexpressing HaCaT cells compared with mock controls (n = 83,779 DARs). (**d–g)** Genome browser tracks showing genomic regions of chromatin accessibility commonly increased in both Arl13B- and GPR161-overexpression HaCaT cells (common DARs) compared to mock controls. Representative inflammatory gene promoters including *IRF7* (**d**), *IKBKE* (**e**), *MAP3K11* (**f**), and *FOSL1* (**g**). Signals were displayed as normalized read counts (y-axis) across genomic coordinates (x-axis). **(h)** HOMER-known motif enrichment analysis of DARs commonly increased in both ARL13B- and GPR161-overexpressing HaCaT cells compared to mock controls. PWM, position–weight matrix. **(i)** Kaplan–Meier analysis of the biologic-free probability after phototherapy in patients with psoriasis, stratified by the number of residual ciliated KCs in Pso-post. Patients were divided into high (≥15 cells/mm) and low (<15 cells/mm) groups based on receiver operating characteristic– derived cut-off values. The shaded areas indicate 95% confidence intervals. **(j)** Waterfall plot showing the number of residual ciliated KCs and subsequent treatments. Patients were ordered according to the number of residual ciliated KCs. Colors indicate subsequent treatment: biologics (red for ≥15 cells/mm; blue for <15 cells/mm) and non-biologic treatment (light red for ≥15 cells/mm; light blue for <15 cells/mm). **(k, l)** Quantification of ciliated cells in HaCaT cells (k) and RPE1 cells (l) following IL-31 stimulation in the presence or absence of SCH. Data are presented as fold change relative to the untreated control condition. Statistical significance was assessed using one-way ANOVA, followed by Dunnett’s multiple comparison test versus the IL-31 group. *P < 0.05, **P < 0.01, **P < 0.0001, \*\*\*\**P* < 0.0001.

Commonly accessible promoters between Arl13B- and GPR161-overexpressing cells included *IRF7*, *IKBKE*, *MAP3K11*, and *FOSL1* (Figure 6d–g), which regulate NF-κB and MAPK signaling. While Arl13B-associated changes were primarily linked to these pathways, GPR161 induced broader remodeling encompassing cytokines and chemokine signaling (Figure S6c and d; see Supporting information). To predict transcription factors mediating these changes, we performed HOMER motif analysis on 569 common DARs between ARL13B- and GPR161-overexpressing cells. Notably, AP-1 family motifs were strongly enriched, with nine of the top ten motifs corresponding to *FOS*, *FOSL*, and *JUN* family members (Figure 6h). Consistently, FOSL1 promoter accessibility increased in both ARL13B- and GPR161-overexpressing cells (Figure 6g). Together, these findings indicate that primary cilia promote chromatin remodeling at inflammatory gene loci, linked to AP-1 associated transcriptional programs downstream of MAPK signaling. This is consistent with the sustained activation of MAPK-related pathways and provides a mechanistic basis for transcriptional priming in ciliated KCs.

### Persistent ciliated KCs after phototherapy are associated with subsequent treatment escalation

To assess the clinical relevance of persistent ciliated KCs, we examined whether their abundance in Pso-post samples was associated with subsequent treatment escalation (Table S7; see Supporting information). Patients with high residual ciliated KCs (≥15) showed a trend toward earlier biologic initiation, although this was not statistically significant (log-rank test, *P* = 0.46) (Figure 6i). Patients who later required biologic therapy also had higher residual ciliated KC counts (Figure 6j). Univariate Cox analysis further supported that high residual ciliated KCs were associated with a 4.37-fold higher risk of subsequent biological therapy, whereas baseline PASI score showed a weaker association (hazard ratio = 1.42), and age and sex were not significant predictors (Table 1). These finding suggest that residual ciliated KCs may reflect persistent disease activity not fully captured by clinical severity scores.

**Table 1.** Univariable Cox proportional hazards analysis for subsequent biologic therapy in psoriasis patients.

| Characteristic | Category | Event/ total | Hazard ratio (95% CI) | P value | Effect size |
|---|---|---|---|---|---|
| Ciliated KCs in pso-post | Cilia ≥15 | 5/7 | 4.37 (0.48–39.56) | 0.01 | NNH 5 |
|  | Cilia <15 | 1/13 |  |  |  |
| PASI | PASI ≥10 | 4/14 | 1.42 (0.15–13.72) | 1 | NNH 8 |
|  | PASI <10 | 1/4 |  |  |  |
| Age | Age ≥70 | 2/10 | 0.72 (0.12–4.37) | 0.63 | NNH 28 |
|  | Age <70 | 4/10 |  |  |  |
| Sex | Male | 3/12 | 1.15 (0.23–5.75) | 0.64 | NNH 1.57 |
|  | Female | 3/8 |  |  |  |

### ERK1/2 inhibitor suppresses IL-31-induced primary cilia formation

Th17 and Th2 cytokines promote primary cilia formation in primary KCs (Figure S7a; see Supporting information).^11^ To identify downstream pathways involved in this process, we screened inhibitors targeting cytokine-induced signaling pathways. Among the inhibitors tested, ERK1/2 inhibition selectively suppressed primary cilia formation (Figure 6k and Figure S7b; see Supporting information), whereas inhibition of other pathways, including STAT signaling, had no significant effect (Figure S7e; see Supporting information). Consistently, phosphorylated ERK1/2 was increased in AD and Pso epidermis, particularly near the basal layer, compared with healthy epidermis (Figure S7d; see Supporting information). These results suggest that ERK1/2 signaling contributes to inflammation-associated primary cilia formation in KCs, and may represent a targetable pathway in AD and psoriasis.

## DISCUSSION

In this study, we identified ciliated KCs as an inflammatory epithelial cell state in AD and Pso. These cells showed enhanced NF-κB and p38 MAPK signaling, increased chromatin accessibility at inflammatory gene loci, and enrichment of AP-1 motifs. Because AP-1 cooperates with NF-κB to amplify inflammatory gene expression, ciliated KCs may acquire a chromatin-primed inflammatory state that contributes to disease chronicity.^48–50^ However, whether the inflammatory stimulation–primary cilia–AP-1 axis directly establishes inflammatory memory remains to be addressed.

Ciliated KCs also showed disease- and environment-dependent programs. In Pso-pre lesions, basal-layer ciliated KCs were enriched for Wnt, Notch, GPCR, and chemokine-related pathways, suggesting a repair-like inflammatory state involving proliferation, early differentiation, and inflammatory niche formation.^51, 52^ Together with the reduced Notch signaling typically linked to suprabasal differentiation defects in inflammatory epidermis, these findings support a biphasic model: basal ciliated KCs adopt a repair-like inflammatory program, whereas suprabasal layers show impaired differentiation. Such heterogeneity indicates that epidermal abnormalities arise from distinct cellular states rather than a uniform defect in differentiation.

Functionally, enhanced primary cilia formation in KCs increased basal MAPK and NF-κB signaling and was associated with AP-1 motif-containing inflammatory chromatin accessibility. GPR161-overexpressing cells showed broader chromatin remodeling than ARL13B-overexpressing cells, possibly reflecting cilia-associated cAMP/PKA signaling.^47, 53^ These findings link primary cilia formation to inflammatory signal amplification and epigenomic priming in KCs.

Clinically, residual ciliated KCs after phototherapy were associated with subsequent treatment escalation, suggesting that they may reflect persistent disease activity not captured by clinical severity alone. The suppression of primary cilia formation by ERK1/2 inhibition suggests that inflammation-associated ciliogenesis may be modifiable, although whether this can be therapeutically exploited remains unclear. Therapeutic targeting of primary cilia should be approached cautiously because systemic cilia defects are associated with immune-related disorders.^54^ Future studies should determine whether KC-specific modulation of primary cilia can suppress inflammatory persistence. In summary, ciliated KCs may contribute to inflammatory memory, disease chronicity, and recurrence in AD and Pso, and may serve as biomarkers of residual disease activity and prognosis.

## ETHICS STATEMENT

This study was performed in accordance with the Declaration of Helsinki. Letter of approval is available on request. Collection of human tissue samples for this study was approved as part of the study protocol. This human study was approved by the Human Research Ethics Committees of the Nagoya City University Clinical Trial Management Center - approval: 60-18-0003, 70-23-0014 and the University of Osaka - approval: yakuhito2019-01, 2019-01-2, 2024-4. All adult participants provided written informed consent to participate in this study.

## DATA AVAILABILITY STATEMENT

The ST data (accession no. GSE327607) and Bulk ATAC-seq data (accession no. GSE328382) are deposited in GEO. The analysis codes are available on the GitHub repository (github.com/Advancedcosmeticscience/ciliated-keratinocyte-analysis).

## CONFLICT OF INTEREST

F.F. was employed by Mandom Corporation. M.I., H.H., A.N., F.F., and M.T. were affiliated with a joint research laboratory established by Mandom Corporation. M.T. was supported by a grant from The Naito Foundation. A.M. has received research grants, consultancy fees, and/or speaker’s fees from AbbVie, Amgen, Boehringer-Ingelheim, Bristol-Myers Squibb, Eli Lilly Japan, Johnson &Johnson, Kyowa Kirin, LEO Pharma, Maruho, Sun Pharma Japan, Torii Pharmaceutical, UCB Japan, and Ushio. All other authors declare no conflict of interest.

## ACKNOWLEDGEMENTS

We thank all the patients at Nagoya City University, Nagoya, Japan. The authors are grateful to Akiko Nishioka, and Yanshu Yang for their helpful assistance with the experiments and discussions. This work was supported by grants from Japan Society for the Promotion of Science Grants-in-Aid for Scientific Research (KAKENHI) (grant nos. JP24K11450 to M.T.), The Naito Foundation (to M.T.), and Osaka University Next-Generation Research Support Program (to M.I. and H.H).

## AUTHOR CONTRIBUTIONS

Conceptualization: M.I., M.T., M.N., A.M.

Methodology: M.I., M.T.

Investigation: M.I., H.H.

Formal analysis: M.I., M.T., Y.H., D.M.

Visualization: M.I.

Funding acquisition: M.I., M.T., F.F. Supervision: K.J.I., M.T., A.M.

Writing—original draft: M.I.

Writing—review and editing: K.J.I., M.T., A.M.

---

**Supplemental Figure 1. Spatial characterization of ciliated cells in ST samples.** (**a**) Line plots show epidermal thickness (left) and PASI scores (right) before and after treatment in paired psoriatic samples. Statistical significance was assessed using paired *t*-test. \**P* < 0.05, \*\*\**P* < 0.001. (**b**) Representative immunofluorescence images of primary cilia in Visum HD sections. Primary cilia were labeled with acetylated α-tubulin (green), basal bodies with pericentrin (red), and nuclei with Hoechst 33342 (blue). White arrows indicate primary cilia. Scale bar: 50 μm (left) and 10 μm (right). (**c**) Immunofluorescence-based spatial distribution of ciliated cells in serial sections corresponding to those used for ST analysis. Three consecutive sections were prepared, with the central section used for the ST and the adjacent sections as shown here. Immunofluorescence staining was performed on all the sections. Representative upper (left) and lower (right) sections are shown, with ciliated cells in red. (**d**) Dot plot showing canonical marker genes used for cell-type annotation. (**e**) Spatial mapping of annotated cell types in Pso-pre.

**Supplemental Figure 2. Characterization of ciliated KCs across inflammatory skin diseases.** (**a**) Spatial prediction of KC layer status in ST sections from healthy, AD, Pso-pre, and Pso-post skin. The basal, spinous, and granular states were defined based on layer-specific gene expression signatures. Prediction scores were overlaid on the tissue sections, with higher scores indicating stronger assignments to each layer. (**b**) Spatial distribution of ciliated and non-ciliated cells in ST sections from AD, Pso-pre-, and Pso-post-skin. Ciliated and non-ciliated cells are highlighted as indicated, respectively. (**c**) Correlation of gene-wise effect sizes (log2 fold change, ciliated versus non-ciliated) between disease conditions restricted to genes that were significantly differentially expressed. (AD vs Pso pre: r = –0.0347, *P* = 6.89 × 10⁻¹; AD vs Pso-post: r = 0.0634, *P* = 4.86 × 10⁻⁵; Pso pre vs Pso post: r = −0.0053, *P* = 9.46 × 10⁻¹)

**Supplemental Figure 3. Ligand-receptor interaction analysis between ciliated KCs and distal cells in psoriatic skin.** Circos plot illustrating ligand-receptor interactions between ciliated KCs and distal (‘far’) cell types in Pso-pre (the two panels on the left) and Pso-post (the two panels on the right) samples. Under each condition, the first and third panels show interactions originating from ciliated KCs (ligands specifically expressed by ciliated KCs), whereas the second and fourth panels show interactions received by ciliated KCs (ligands derived from non-ciliated KCs, FBs, and ECs interacting with receptors preferentially expressed in ciliated KCs).

**Supplemental Figure 4. Ciliated KCs exhibit distinct inflammatory signaling and spatial interactions in AD.** (**a**) GO analysis of DEGs between ciliated and non-ciliated KCs in AD samples. (**b**) Bar plot showing significantly enriched signaling pathways identified by GSEA of genes upregulated in ciliated versus non-ciliated KCs in AD. (**c**) Circos plot illustrating selected signaling pathways (MAPK–AP-1, RPTP, and ROBO) enriched in ciliated KCs relative to non-ciliated KCs in AD, based on the GSEA results shown in (**b**) and their associated DEGs. (**d**) Heatmaps showing the number of ligand–receptor pairs between ciliated KCs and adjacent and far cells in AD samples. (**e**) Circos plot illustrating ligand–receptor interactions originating from ciliated KCs in AD. Ligands preferentially expressed by ciliated KCs are predicted to interact with receptors on adjacent non-ciliated KCs, FBs, ECs, and nerve cells. (**f**) Circos plot illustrating ligand– receptor interactions in ciliated KCs in AD. Ligands derived from adjacent non-ciliated KCs, ECs, mast cells (Mast), and melanocytes (MLNC) are predicted to interact with receptors preferentially expressed by ciliated KCs.

**Supplemental Figure 5. Distinct and shared chromatin accessibility programs induced by ARL13B and GPR161.** (**a**) Quantification of p38 MAPK activation shown in Fig. 5C, expressed as the ratio of p-p38 MAPK to total p38 MAPK (p-p38/total p38). Statistical significance was assessed by one-way ANOVA followed by Dunnett’s multiple comparison test versus the control group. (**b**) Quantification of p-IκBα shown in Figure 5c, expressed as the ratio of p-IκBα to total IκBα (p-IκBα/total IκBα). Statistical significance was assessed by one-way ANOVA followed by Dunnett’s multiple comparison test versus the control group. \*\*\**P* < 0.001.

**Supplemental Figure 6. Distinct and shared chromatin accessibility programs induced by ARL13B and GPR161.** (**a**) Principal PCA of ATAC-seq chromatin accessibility profiles across mock-, ARL13B-, and GPR161-overexpressing HaCaT cells. (**b**) Venn diagram showing the overlap of DARs located in the promoter regions of ARL13B- and GPR161-overexpressing HaCaT cells. (**c**) Replicate-pooled ATAC-seq peak tracks showing chromatin accessibility at promoter regions of genes associated with NF-κB (*RELA*, *TRAF6*, *TIRAP*, *PELI3*) and MAPK (*MAP2K1*, *MAP2K5*, *PAK1*, *FYN*) signaling pathways in ARL13B-overexpressing HaCaT cells. Gene orientation is indicated by + (forward strand) and − (reverse strand). (**d**) Replicate-pooled ATAC-seq peak tracks showing chromatin accessibility at promoter regions of genes associated with NF-κB (*IKBKB*, *TRAF2*, *CARD11*, *TNIP1*), MAPK (*MAP3K8*, *MAP3K13*, *MKNK1*, *FOS*), TLR (*TLR2*, *TLR4*, *CD14*, *TICAM1*) signaling pathways, and cytokines/chemokines (*CXCL14*, *CXCL8*, *TNFSF10*, *IL1B*) in GPR161-overexpressing HaCaT cells. Gene orientation is indicated by + (forward strand) and − (reverse strand).

**Supplemental Figure 7. ERK1/2 signaling promotes cytokine-induced primary cilia formation. (a)** Quantification of ciliated cells after treatment with IL-31, STAT inhibitors, or IL-31 combined with each STAT inhibitor. Data are presented as fold change relative to the untreated control. Statistical significance was assessed using one-way ANOVA followed by Dunnett’s multiple comparison test versus the IL-31 group. **(b)** Immunofluorescence images of p-Erk and primary cilia. Ciliary axonemes were labeled with acetylated α-tubulin (green), centrosomes with pericentrin (red), p-Erk with anti-p-ERK antibody (black), and nuclei with Hoechst 33342 (blue). White arrows indicate primary cilia. Scale bar: 15 µm.

## SUPPLEMENTARY MATERIALS AND METHODS

### Study design

We employed high-resolution spatial transcriptomics (ST) in conjunction with immunofluorescence staining to characterize ciliated cells in human inflammatory skin diseases including AD and Pso. Skin biopsy specimens were obtained from 7 HCs, 15 patients with AD, and 20 patients with Pso. Pso-paired epidermal samples were collected from the same patients before and after phototherapy to assess the treatment-associated changes. Primary cilia were identified in tissue sections using ciliary axoneme and centrosomal markers and subjected to ST analysis. This approach enabled the integration of the spatial localization of ciliated cells with single-cell–resolved whole-transcriptome profiles while preserving the intact tissue architecture. The transcriptomic features of ciliated cells were analyzed across disease states, treatment conditions, and HCs with a specific focus on epidermal KCs. Additionally, spatial proximity analyses were conducted to analyze local cellular associations surrounding the ciliated cells. We performed in vitro experiments, including ATAC-seq, to investigate the relationship between inflammatory memory and primary cilia.

### Human samples and IRB statement

All experiments were approved by the Human Research Ethics Committees of the Nagoya City University Clinical Trial Management Center (No. 60-18-0003, 70-23-0014) and the University of Osaka (yakuhito2019-01, 2019-01-2, 2024-4). This study was conducted in accordance with the Declaration of Helsinki Principles and ethical guidelines for medical and biological research involving human subjects in Japan, and informed consent was obtained from all adult participants. Archived FFPE skin samples from patients with Pso and AD treated at Nagoya City University Hospital were used in this study. After preparation of 3 µm-thick tissue sections, the slides were transported to the University of Osaka for ST analysis. Clinical assessments and diagnoses were performed by experienced dermatologists and standardized through regular consensus meetings within the institution. Patient characteristics and clinical information (Supplementary Table S1 and S7).

### RNA quality assessment

RNA quality was assessed by measuring the percentage of RNA fragments longer than 200 base pairs (DV200) using 5 μm-thick sections cut from each FFPE tissue block. DV200 was measured using an Agilent 2100 Bioanalyzer with a Eukaryote Total RNA Pico Series II Kit. Samples with DV200 values greater than 30% were used for ST analysis.

### Tissue preparation and primary cilia detection by immunostaining

ST analysis was performed using the 10x Genomics Visium HD Spatial Gene Expression platform on FFPE human adult skin tissue samples. The dataset included paired pre- and post-treatment psoriatic skin samples obtained from the same seven patients (n = 7 Pso-pre and n = 7 Pso-post), AD samples (n = 10), and HC skin samples (n = 5).

Three consecutive tissue sections with thicknesses of 6, 3, and 6 μm were prepared, and the central 3 μm-thick section was used for ST analysis. The primary cilia and nuclei were identified by immunofluorescence staining in all three consecutive sections. The upper and lower sections were processed using standard immunofluorescence staining protocols, whereas the central sections were processed according to the manufacturer’s instructions for ST. The spatial positions of the primary cilia and nuclei in the upper and lower sections were manually aligned with those in the central section to identify ciliated cells within the ST section. Immunostained tissue sections were mounted onto ST analysis areas following the manufacturer’s protocol for FFPE samples.

Three serial sections were prepared from the FFPE tissue blocks. The upper and lower 6 µm sections were collected for standard immunostaining, and the middle 3 µm section was collected for ST analysis. To remove wrinkles, the section for the ST analysis was floated on a 42°C water bath and then transferred to a room-temperature water bath to allow the paraffin to solidify. Tissue sections were trimmed to approximately 1 mm × 1 mm using fine scissors and forceps in a room-temperature water bath. The trimmed tissue section was placed within the 6.5 mm × 6.5 mm capture area of a SuperFrost Plus slide (Epredia, J1800AMNZ). After placing each sample, the slide was dried at 42°C for at least 2 min to prevent tissue detachment. Slides with mounted tissue sections were stored at 4°C until deparaffinization was performed. The slides were incubated on a section hot plate at 60°C for 2 h to melt the paraffin and then cooled to room temperature for 5 min. Slides were deparaffinized by sequential placement in xylene and ethanol. Slides were assembled into a tissue slide cassette, and a de-crosslinking Mix (10x Genomics, PN-1000668) was added to fully cover the tissue section. After de-crosslinking, slides were gently rinsed with phosphate-buffered saline (PBS). Slides were blocked with a blocking buffer consisting of 1× Block and Stain Buffer (Visium FFPE Reagent Kit, 10x Genomics) supplemented with an RNase inhibitor (Protector RNase Inhibitor, Roche) for 30 min at room temperature. Primary antibodies diluted in an antibody solution containing bovine serum albumin (BSA) (Thermo Fisher Scientific, AM2616), an RNase inhibitor (Roche, 03335399001), and Tween-20 in PBS were applied to the tissue sections and incubated for 1 h at room temperature. The following antibodies and dilutions were used: mouse anti-acetylated tubulin (Sigma, T6793, 1:25) and rabbit anti-pericentrin/kendrin (Thermo Fisher Scientific, IHC-00264, 1:50). The sections were incubated with secondary antibodies and Hoechst 33342 (H3570, 1:500; Thermo Fisher Scientific) for 1 h at room temperature. The secondary antibodies included donkey anti-mouse IgG Alexa Fluor 488 (A21202, 1:500; Thermo Fisher Scientific) and donkey anti-rabbit IgG Alexa Fluor 594 (A21207, 1:500; Thermo Fisher Scientific) diluted in nuclease-free water. After washing with PBS, the sections were mounted using SlowFade Diamond Antifade Mountant (Thermo Fisher Scientific, S36963) supplemented with glycerol (Sigma-Aldrich, 12-1120-5) and coverslipped. Fluorescence images were acquired using a slide scanner (VS200; Evident Scientific). After fluorescence imaging, the coverslips were removed by immersing the slides in 1× PBS until they detached naturally without mechanical force. The slides were then rinsed three times in PBS to remove the residual mounting medium, air-dried for 5 min, and placed in new tissue slide cassettes. Slides were immediately processed using the CytAssist workflow according to the manufacturer’s instructions.

### ST data analysis

Sequencing and pre-processing libraries were sequenced on an MGI DNBSEC-G400 platform, achieving a minimum depth of 400 million reads per slide (approximately 18 million reads per sample). Raw sequencing data were processed using Space Ranger (v4.0.1) with the Visium Human Transcriptome Probe Set v2.0 and the GRCh38-2020-A reference genome.

### Cell segmentation and cell type annotation

Cell segmentation was performed using StarDist on the Hoechst-stained nuclear channel of the immunofluorescence images. Transcript counts from 2 μm bins were aggregated into a single-cell profile. Briefly, the algorithm assigns weights to each 2 μm bin based on the StarDist masks. To capture extranuclear transcripts, we applied Gaussian blurring by assigning reduced distance-dependent weights to counts outside the primary segment. Consequently, the final cell-by-feature matrix contained fractional UMI counts. To eliminate misidentified cells and segmentation artifacts, we subsequently filtered out cells with a count density below 0.5 UMIs per 2 μm bin. Cell types were annotated using a custom automated classification tool based on marker gene expression. The reference marker genes were derived from previously published human skin datasets.^1^ This algorithm classifies query cells by calculating the cosine similarity between the reference gene expression vector of a specific cell type and the expression vector of the query cell. The accuracy of the automated annotations was manually evaluated by dermatology experts and demonstrated an overall good agreement with manual histological and marker gene-based annotations.

### Sub-clustering of epidermal KCs

Clusters were obtained using the FindNeighbors and FindClusters functions, with the resolution set at 0.5. Cluster marker genes were identified using the FindAllMarker function. Cell types were annotated using overlapping cluster markers with canonical cell-type signature genes.

### Differential expression and pathway analysis

Gene set enrichment analysis was performed to identify the biological pathways specifically associated with ciliated KCs in the Pso and AD epidermis. DEGs between ciliated and non-ciliated KCs in the basal layer were computed from ST data using Seurat with the Wilcoxon rank-sum test. Cells with low feature counts (nFeature_RNA < 10) were excluded from the analysis. Gene expression values were log-normalized.

For Gene Set Enrichment Analysis (GSEA), the genes were ranked based on their average log2 fold change. GO Biological Process (GO: BP) gene sets were obtained from the Molecular Signatures Database (MSigDB) using the msigdbr package (collection C5, GO: BP, *Homo sapiens*). Reactome gene sets were obtained using the msigdbr package (*Homo sapiens*; collection C2; sub-collection CP: REACTOME). Statistical significance was set at *P* < 0.05. overexpressed.

### Ligand–receptor interaction analysis

Ligand–receptor interaction analysis was performed using CellPhoneDB (v5.0.0) to investigate the spatial cell–cell communication networks associated with ciliated KCs.^2^ Spatial coordinates obtained from the ST data were used to define neighboring cell populations. Cells located within 50 μm of ciliated KCs were defined as adjacent cells, whereas cells located between 50–100 μm were defined as distal cells. The distances were calculated based on the Euclidean distance between spatial coordinates.

Ligand–receptor interactions between ciliated KCs and adjacent/distant cells were inferred using CellPhoneDB statistical analysis with permutation testing. Pairs with a p-value greater than 0.05 were excluded from further analysis. In parallel, ligand–receptor interactions were computed between non-ciliated KCs and adjacent/distant cells.

To identify the signaling pathways specifically received or secreted by ciliated KCs, the interaction results derived from non-ciliated KCs were subtracted from those derived from ciliated KCs. Interaction pairs were matched by interaction pair identity and receiver cell type, and the interaction scores were compared. Cilia-specific ligand–receptor candidates were defined as interaction pairs detected in ciliated KCs, but absent in non-ciliated KCs.

### Cell culture

HaCaT cells and Retinal Pigment Epithelium 1 (RPE1) cells were cultured in Dulbecco’s modified Eagle’s medium (DMEM; Gibco, C1199500) containing 20% fetal bovine serum (Gibco, 10270-106), and a 1× penicillin–streptomycin–amphotericin B suspension (FUJIFILM Wako, 161-23181).

Stable HaCaT cell lines overexpressing ARL13B or GPR161 were established by lentiviral transduction using pLVX-EGFP-ARL13B-puro or pLVX-GFP-GPR161-puro vectors. Puromycin selection (2 µg/ml) was initiated 3 days after infection and maintained for 3 weeks. Single cells were isolated and expanded to establish the clonal cell lines.

### Cytokine and inhibitor treatments

HaCaT and RPE1 cells were maintained in DMEM supplemented with 10% fetal bovine serum and 1% antibiotic–antimycotic at 37°C in 5% CO₂. RPE1 cells were seeded in 24-well plates at 5.0 × 10³ cells per well and stimulated with IL-13, IL-31, IL-17A, or TNF-α at 10 ng/ml for 24 h. HaCaT cells were seeded at 2.0 × 10⁵ cells per well and stimulated with IL-13, IL-31, or IL-17A at 10 ng/ml for 72 h.

For inhibitor experiments, cells were pretreated with pathway inhibitors for 1 h before stimulation with IL-31. The inhibitors used were WP1066 for STAT3 (10nM; Selleck, S2796), Fludarabine for STAT1 (10nM; Selleck, S1491), STAT5-IN-1 for STAT5 (10nM; Selleck, S6784), AS1514799 for STAT6 (100nM; Selleck, S8685), and SCH772984 for ERK1/2 (2.5nM; Selleck, S7101). After inhibitor pretreatment, cells were stimulated with IL-31 at 10 ng/ml and cultured for 24 h for RPE1 cells or 72 h for HaCaT cells.

### Immunofluorescence staining and quantification

FFPE tissue sections from patients with Pso or AD, and HCs were heated at 60°C for 1 h, deparaffinized, and rehydrated. The slides were immersed in boiled 1× Tris-EDTA buffer (10 mM Tris base and 1 mM EDTA, pH 9.0). After cooling, the slides were blocked for 1 h at room temperature in PBS containing 0.1% TritonX-100 and 10% fetal bovine serum, followed by an overnight incubation at 4 °C with primary antibodies diluted in PBS containing 0.1% Tween 20. The primary antibodies used were mouse anti-acetylated tubulin (T6793, 1:500; Sigma) and rabbit anti-pericentrin/kendrin (IHC-00264, 1:500; Thermo Fisher Scientific). The sections were washed with PBS containing 0.1% Tween 20 before and after incubation with secondary antibodies and Hoechst 33342 (H3570, 1:1000) for 1 h at room temperature. Secondary antibodies included donkey anti-mouse IgG Alexa Fluor 488 (A21202, 1:500; Thermo Fisher Scientific) and donkey anti-rabbit IgG Alexa Fluor 594 (A21207, 1:500; Thermo Fisher Scientific), diluted in PBS containing 0.1% Tween 20. After washing with PBS, the sections were mounted on glass slides using ProLong Glass antifade mountant (Thermo Fisher Scientific, P36980). Fluorescent images were acquired using an Olympus FV3000 confocal microscope (Olympus) and a slide imaging scanner SLIDEVIEW VS200 (Olympus).

### Quantification of primary cilia

The percentage of ciliated cells was quantified on 5 μm-thick skin sections using FV31S-SW software (Olympus). Z-stacks were acquired by defining the upper and lower focal planes with detectable fluorescence using a 100× objective, and the step size was determined using system-optimized settings. Z-stack images were processed to generate maximum intensity projections. The percentage of ciliated cells was calculated as the number of cilia positive for both acetylated tubulin and pericentrin and normalized to the horizontal tissue length per field of view.

### Protein extraction and western blot analysis

Protein extracts were prepared by lysing the cells for 30 min in RIPA buffer (Santa Cruz Biotechnology, sc-24948) containing protease (Roche, 0469159001) and phosphatase inhibitors (Roche, 04906837001). The proteins were denatured by adding Laemmli sample buffer (Bio-Rad) and heating at 95°C for 5 min. Protein samples were separated on a sodium dodecyl sulfate–polyacrylamide gel (Bio-Rad, Hercules, CA, USA; 16101173) and transferred onto polyvinylidene difluoride membranes. Membranes were blocked with Blocking Reagent N102 (NOF Corporation, S410-03022) and incubated overnight 4°C with primary antibodies: phosphorylated p38 (CST, 4511, 1:1000), p38 (CST, 8690, 1:1000), phosphorylated IκBα (CST, 9246, 1:1000), and IκBα (CST, 4814, 1:1000). After incubation with secondary antibodies, the protein bands were visualized using an Amersham Imager 600 (GE Healthcare) and quantified using ImageJ software.

### Bulk ATAC-seq library preparation

ATAC-seq libraries were prepared from HaCaT cells by using ATAC-seq Kit (Active Motif, Inc., 53150) according to the manufacturer’s instructions. All reagents used in the following procedures were provided in the kit unless otherwise specified. For nuclear isolation, 50,000–100,000 cells were centrifuged at 500 × *g* for 5 min at 4°C, then washed once with PBS and centrifuged again at 500 × *g* for 5 min. The supernatant was removed, and cells were resuspended in ATAC lysis buffer. Nuclei were pelleted by centrifugation at 500 × *g* for 10 min at 4°C. The nuclei pellet was resuspended in the transposition reaction mix and incubated at 37°C for 30 min with shaking. Following transposition, DNA was purified using a silica column–based purification method, and purified DNA was amplified using indexed primers and polymerase. PCR was performed under the following conditions: 72°C for 5 min, 98°C for 30 s, followed by 10 cycles of 98°C for 10 s, 63°C for 30 s, and 72°C for 1 min. Libraries were purified using SPRI beads according to the manufacturer’s instructions and quantified before sequencing.

### Bulk ATAC-seq processing and reading alignment

Paired-end bulk ATAC-seq libraries were processed using a standardized pipeline. Adapter and low-quality bases were removed from raw FASTQ files prior to alignment. Reads were aligned to the human reference genome (hg38) using Bowtie2, with parameters optimized for ATAC-seq. Alignments were converted to a binary alignment map (BAM) format, coordinate-sorted, and indexed using SAMtools. PCR duplicates were removed to generate deduplicated BAM files for downstream quantification and differential accessibility analysis.

### Bulk ATAC-seq peak calling

Accessible chromatin regions were identified from deduplicated BAM files using MACS3 peak calling with the appropriate ATAC-seq settings, generating narrow peak outputs for each sample. For downstream differential analysis, peak coordinates were harmonized across all samples via DiffBind’s consensus peak set construction and summit recentering (summits = 250 bp), ensuring a common set of genomic intervals for read counting and statistical testing.

### Bulk ATAC-seq differential chromatin accessibility analysis (DiffBind/DESeq2)

Differential accessibility was assessed using DiffBind with DESeq2 normalization and modeling. Read counts were computed over the consensus peak set from deduplicated coordinate-sorted BAM files, treating the reads as paired-end fragments. Peaks overlapping the blacklisted ENCODE regions were removed before analysis. Differential testing was performed for the contrasts mock vs. ARL13B and mock vs. GPR161, and multiple testing corrections were applied using the Benjamini–Hochberg procedure. Peaks were considered significantly differentially accessible at a false discovery rate (FDR) < 0.05.

### Identification of concordantly opened promoter regions in ARL13B and GPR161

To identify promoter regions selectively opened in both ARL13B and GPR161 relative to mock, promoter intervals were defined as ±500 bp around annotated transcription start sites using TxDb.Hsapiens.UCSC.hg38.knownGene. Consensus peaks were classified as promoter overlapping if they intersected the promoter intervals (ignored strands). The directionality of accessibility changes was defined using normalized peak counts; peaks were required to exhibit higher mean normalized accessibility in both ARL13B and GPR161 compared to mock (ARL_mean > mock_mean and GPR_mean > mock_mean). To enrich promoters with low baseline accessibility in mock, mock-low promoters were defined as those with mock mean normalized counts within the bottom 5th percentile across standard chromosomes. High-confidence concordantly opened promoters were defined as those satisfying the following conditions: (i) promoter overlap, (ii) significant differential accessibility in both mock vs. ARL13B and mock vs. GPR161 contrasts (FDR < 0.05), (iii) concordant opening directionality in both mutants relative to mock based on normalized counts, and (iv) low baseline accessibility in mock (bottom 5th percentile).

### Motif enrichment analysis

For pathway-level interpretation, promoter peaks meeting concordant opening criteria were annotated to nearby genes using ChIPseeker with a ±500 bp transcription start site window. Significantly enriched known transcription factor motifs were identified based on Benjamini–Hochberg–adjusted q-values (< 0.05). Position-weight matrices for enriched motifs were obtained from the JASPAR2020 database using the TFBSTools package, and motif sequence logos were generated using the ggseqlogo package. The motifs were grouped into transcription factor families based on their annotations.

### Chromatin accessibility track generation and promoter profiling

Deduplicated, coordinate-sorted, paired-end ATAC-seq BAM files were converted to normalized coverage tracks using deepTools (bamCoverage v3.5.4) with CPM normalization (bin size 10 bp), minimum mapping quality of 30, and fragment extension (extendReads). The promoter regions were defined as transcription start sites annotated in TxDb.HSapiens.UCSC.hg38.knownGene, exported as BED, and profiled using deepTools computeMatrix (reference-point mode; TS38. KnownGeneoup-averaged promoter accessibility profiles for the mock, ARL13B, and GPR161 conditions were visualized using deepTools plotProfile (mean across replicates).

### Visualization and quality control

Normalized coverage tracks were generated from BAM files and visualized using the Integrative Genomics Viewer to inspect representative loci and confirm consistency across biological replicates. Peak reproducibility and differential accessibility patterns were examined visually to ensure biological reproducibility.

### Clinical association analysis

The clinical relevance of residual ciliated keratinocytes (KCs) after phototherapy was assessed in patients with psoriasis with available post-phototherapy samples and follow-up data. Residual ciliated KCs were quantified as the number of ciliated KCs per millimeter of epidermis. Subsequent treatment escalation was defined as initiation of biological therapy after phototherapy.

Receiver operating characteristic analysis identified 15 ciliated KCs/mm as the optimal cutoff for predicting subsequent biological therapy (AUC = 0.786, sensitivity = 83.3%, specificity = 85.7%). Patients were classified into high and low residual ciliated KC groups using this cutoff. Biologic-free survival was analyzed using the Kaplan–Meier method and compared with the log-rank test. Associations between subsequent biological therapy and clinical variables, including age ≥70 years, male sex, baseline PASI ≥10 and residual ciliated KCs ≥15/mm, were evaluated using Fisher’s exact test and univariate Cox proportional hazards regression.

### Data and materials availability

ST data are available in GEO under accession GSE327607. BAM files for bulk ATAC-seq are available at GSE328382. All code used for data processing, analysis, and figure generation is available on GitHub. All data needed to evaluate the conclusions in the paper are present in the paper. All materials used or generated in this study are commercially available or will be supplied upon reasonable request.

### Statistical analysis

Sample sizes were not predetermined based on statistical methods but were chosen according to the standards of the field (at least three independent biological replicates for each condition), which generated a sufficient number of tissues and cells and provided sufficient statistics for the effect sizes of interest. All statistical tests, except for ST and ATAC-seq, were performed using GraphPad Prism 9 or R. For comparisons between two groups, unpaired and paired two-tailed Student’s *t*-tests were used. One-way ANOVA followed by Dunn’s multiple comparison test was used for comparisons between groups. The Benjamini–Hochberg procedure was used to adjust the FDR. Individual values are presented as bar plots. *P* values were considered significant when < 0.05 (\**P* < 0.05, \*\**P* < 0.01, \*\*\**P* < 0.001,\*\*\*\**P* < 0.0001).

## SUPPLEMENTARY TABLES

**Supplemental Table S1.** Patient clinical characteristics for ST.

**Supplemental Table S2.** Table listing differentially expressed genes between ciliated KCs and layer-matched non-ciliated KCs in AD, Pso-pre, and Pso-post. DEGs were identified using pseudobulk aggregation of ST data followed by differential expression analysis. Adjusted P values were calculated using the Benjamini–Hochberg method.

**Supplemental Table S3.** Table listing GO terms enriched in ciliated KCs compared with non-ciliated KCs across AD, Pso-pre, and Pso-post. GO enrichment analysis was performed on differentially expressed genes identified from pseudobulk analysis. The table includes the number of genes, NES, P values, and core-enriched genes for each GO term.

**Supplemental Table S4.** Table listing Reactome pathways enriched in ciliated KCs compared with non-ciliated KCs across AD, Pso-pre, and Pso-post. Reactome pathway analysis was performed on differentially expressed genes identified from pseudobulk analysis of layer-matched cells. The table includes the number of genes, NES, P values, and core-enriched genes for each pathway.

**Supplemental Table S5.** Table listing ligand–receptor interactions involving ciliated KCs across AD, Pso-pre, and Pso-post. Cell–cell communication analysis was performed using CellPhoneDB. Interactions are stratified based on whether ciliated KCs act as sender or receiver cells and further categorized by spatial proximity, with adjacent cells defined as within 50 µm and distal cells defined as 50–100 µm from ciliated KCs. The table includes interacting ligand–receptor pairs, ligand and receptor genes, interacting cell types, and adjusted interaction scores.

**Supplemental Table S6.** Table listing ATAC-seq data from HaCaT cells overexpressing ARL13B or GPR161 compared with mock controls. DARs were identified, and promoter-associated peaks and corresponding genes are provided. The table includes genomic coordinates, gene annotations, and differential accessibility metrics for ARL13B and GPR161 overexpression conditions.

**Supplemental Table S7.** Clinical characteristics and association between ciliated keratinocytes and disease progression in psoriasis.

