## Supplemental Figures for "Ciliated keratinocytes define a spatially restricted pro-inflammatory state in psoriasis and atopic dermatitis"

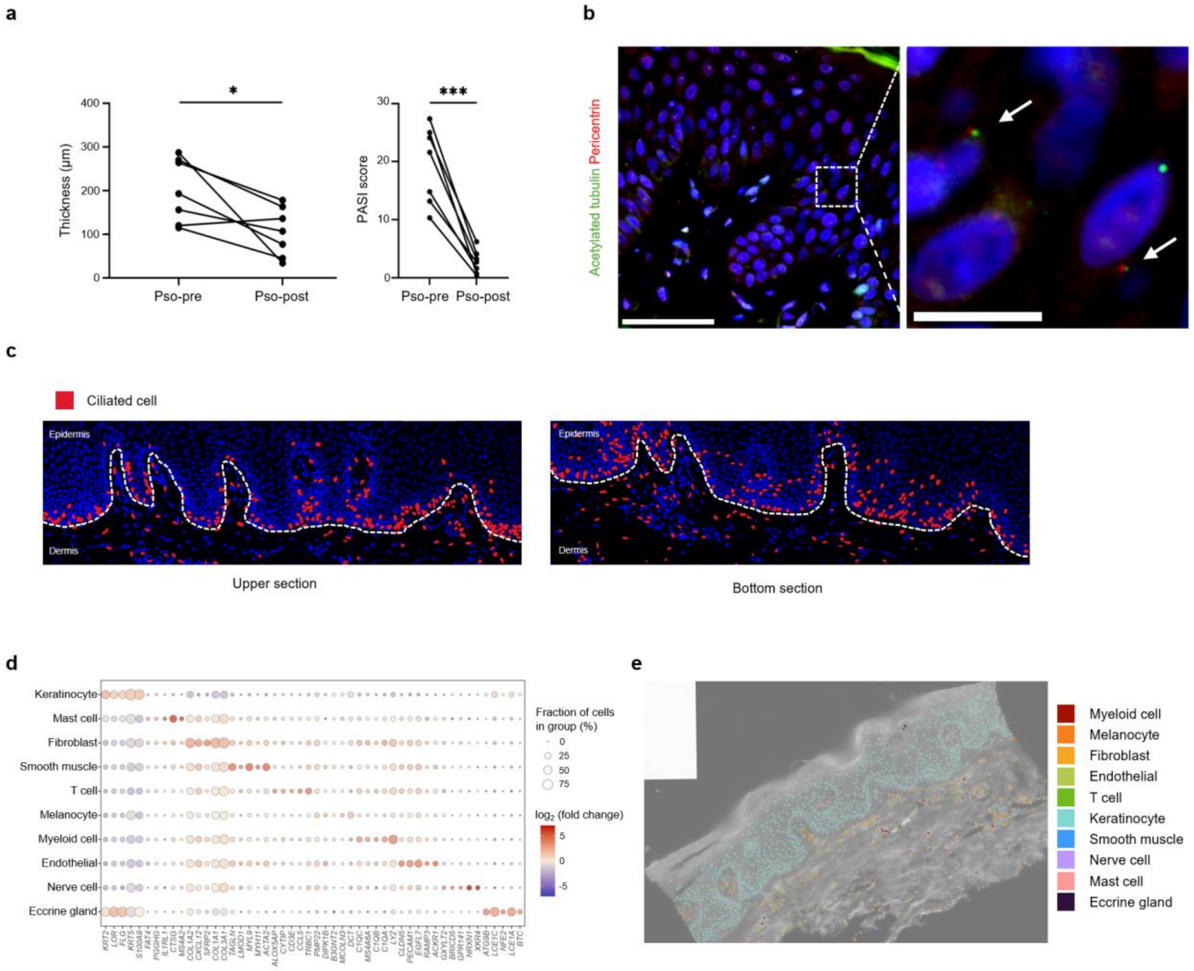


**Supplemental Figure 1. Spatial characterization of ciliated cells in ST samples.** (**a**) Line plots show epidermal thickness (left) and PASI scores (right) before and after treatment in paired psoriatic samples. Statistical significance was assessed using paired *t*-test. *$P < 0.05$, ***$P < 0.001$. (**b**) Representative immunofluorescence images of primary cilia in Visum HD sections. Primary cilia were labeled with acetylated α-tubulin (green), basal bodies with pericentrin (red), and nuclei with Hoechst 33342 (blue). White arrows indicate primary cilia. Scale bar: 50 μm (left) and 10 μm (right). (**c**) Immunofluorescence-based spatial distribution of ciliated cells in serial sections corresponding to those used for ST analysis. Three consecutive sections were prepared, with the central section used for the ST and the adjacent sections as shown here. Immunofluorescence staining was performed on all the sections. Representative upper (left) and lower (right) sections are shown, with ciliated cells in red. (**d**) Dot plot showing canonical marker genes used for cell-type annotation. (**e**) Spatial mapping of annotated cell types in Pso-pre.

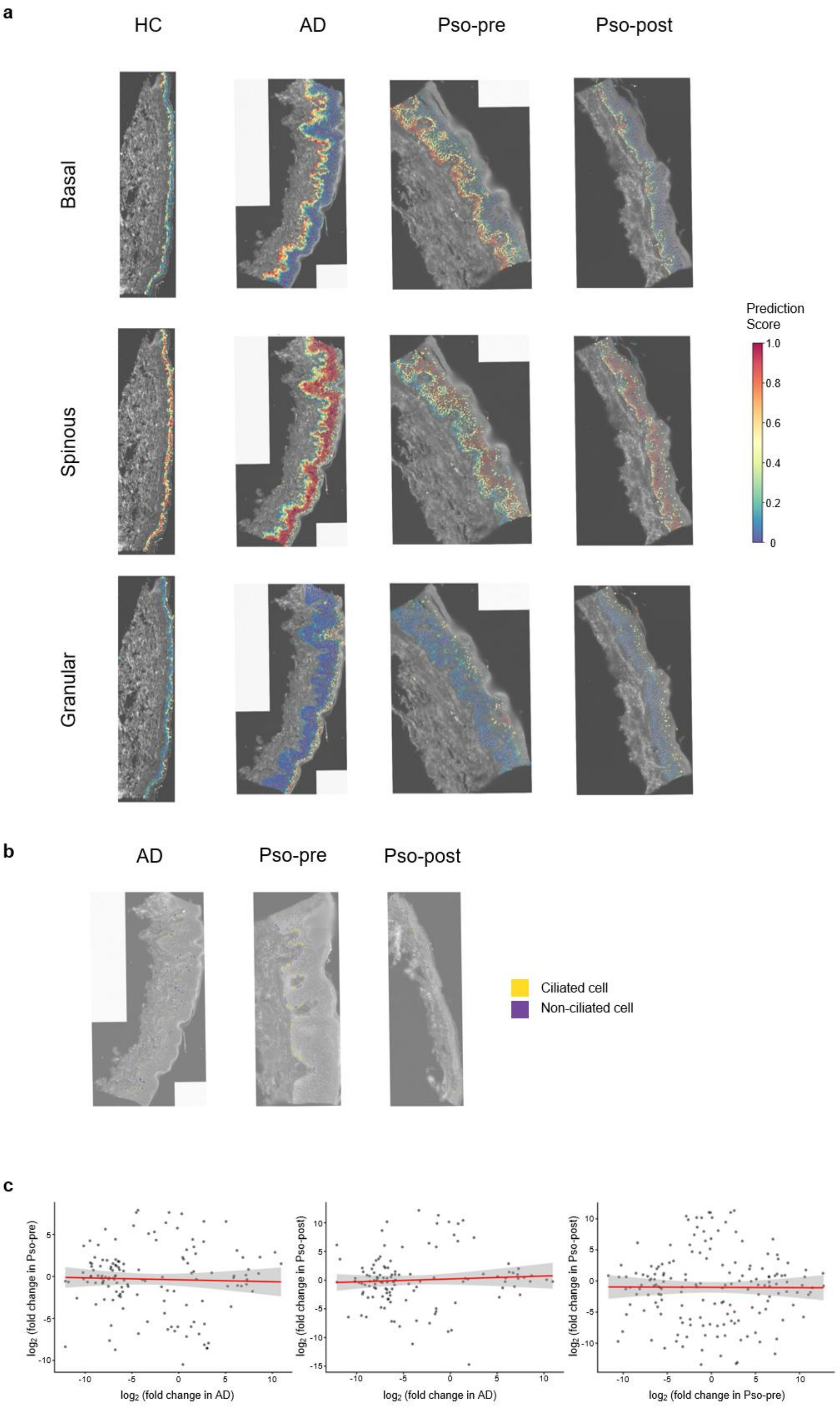
a
HC
AD
Pso-pre
Pso-post
Basal
Spinous
Granular
Prediction Score
1.0
0.8
0.6
0.4
0.2
0
b
AD
Pso-pre
Pso-post
Ciliated cell
Non-ciliated cell
c
log2 (fold change in Pso-pre)
log2 (fold change in AD)
log2 (fold change in Pso-post)
log2 (fold change in AD)
log2 (fold change in Pso-post)
log2 (fold change in Pso-pre)

**Supplemental Figure 2. Characterization of ciliated KCs across inflammatory skin diseases.** (**a**) Spatial prediction of KC layer status in ST sections from healthy, AD, Pso-pre, and Pso-post skin. The basal, spinous, and granular states were defined based on layer-specific gene expression signatures. Prediction scores were overlaid on the tissue sections, with higher scores indicating stronger assignments to each layer. (**b**) Spatial distribution of ciliated and non-ciliated cells in ST sections from AD, Pso-pre-, and Pso-post-skin. Ciliated and non-ciliated cells are highlighted as indicated, respectively. (**c**) Correlation of gene-wise effect sizes (log2 fold change, ciliated versus non-ciliated) between disease conditions restricted to genes that were significantly differentially expressed. (AD vs Pso pre: r = –0.0347, $P = 6.89 \times 10^{-1}$; AD vs Pso-post: r = 0.0634, $P = 4.86 \times 10^{-5}$; Pso pre vs Pso post: r = −0.0053, $P = 9.46 \times 10^{-1}$)

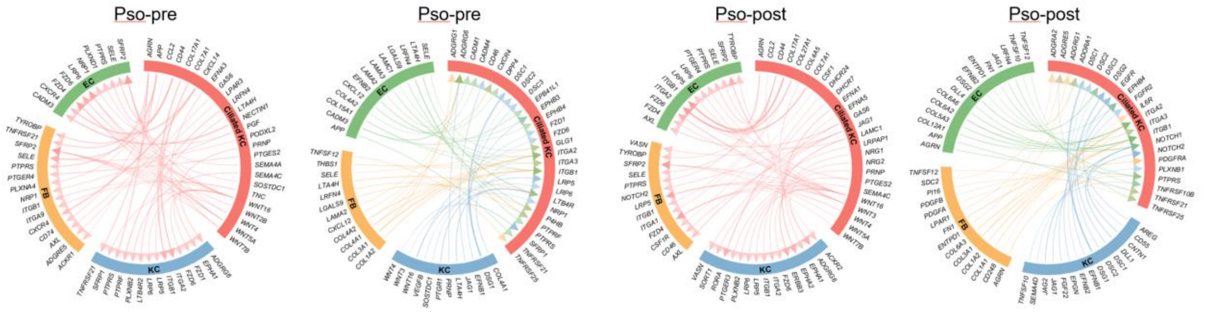


**Supplemental Figure 3. Ligand-receptor interaction analysis between ciliated KCs and distal cells in psoriatic skin.** Circos plot illustrating ligand-receptor interactions between ciliated KCs and distal ('far') cell types in Pso-pre (the two panels on the left) and Pso-post (the two panels on the right) samples. Under each condition, the first and third panels show interactions originating from ciliated KCs (ligands specifically expressed by ciliated KCs), whereas the second and fourth panels show interactions received by ciliated KCs (ligands derived from non-ciliated KCs, FBs, and ECs interacting with receptors preferentially expressed in ciliated KCs).

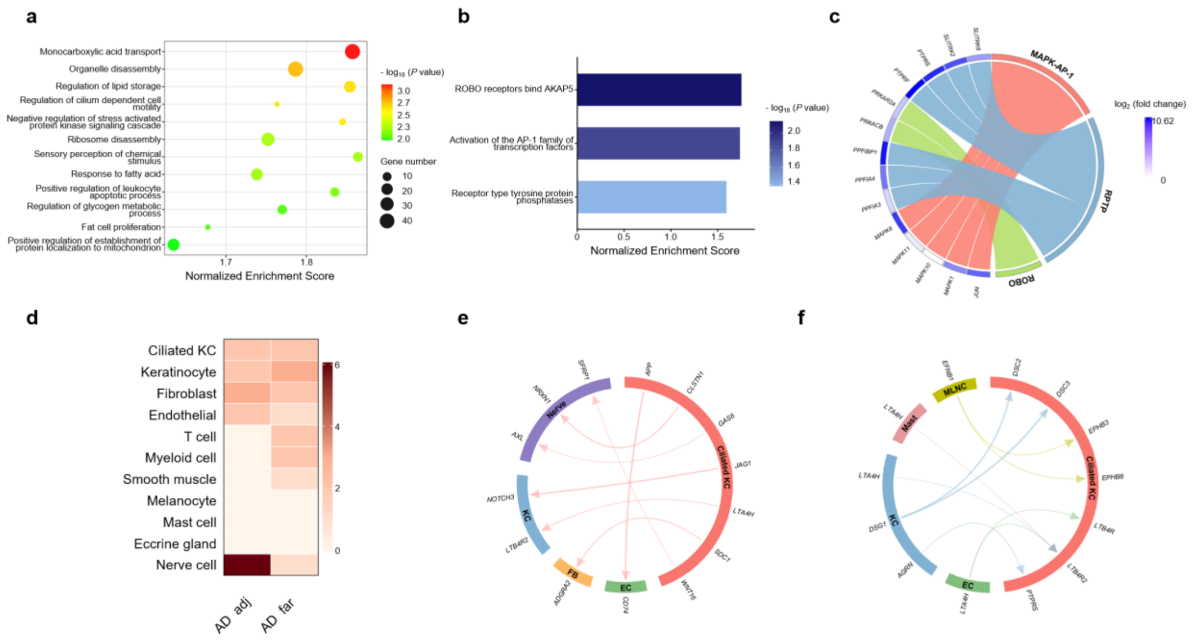


**Supplemental Figure 4. Ciliated KCs exhibit distinct inflammatory signaling and spatial interactions in AD.** (**a**) GO analysis of DEGs between ciliated and non-ciliated KCs in AD samples. (**b**) Bar plot showing significantly enriched signaling pathways identified by GSEA of genes upregulated in ciliated versus non-ciliated KCs in AD. (**c**) Circos plot illustrating selected signaling pathways (MAPK–AP-1, RPTP, and ROBO) enriched in ciliated KCs relative to non-ciliated KCs in AD, based on the GSEA results shown in (**b**) and their associated DEGs. (**d**) Heatmaps showing the number of ligand–receptor pairs between ciliated KCs and adjacent and far cells in AD samples. (**e**) Circos plot illustrating ligand–receptor interactions originating from ciliated KCs in AD. Ligands preferentially expressed by ciliated KCs are predicted to interact with receptors on adjacent non-ciliated KCs, FBs, ECs, and nerve cells. (**f**) Circos plot illustrating ligand–receptor interactions in ciliated KCs in AD. Ligands derived from adjacent non-ciliated KCs, ECs, mast cells (Mast), and melanocytes (MLNC) are predicted to interact with receptors preferentially expressed by ciliated KCs.

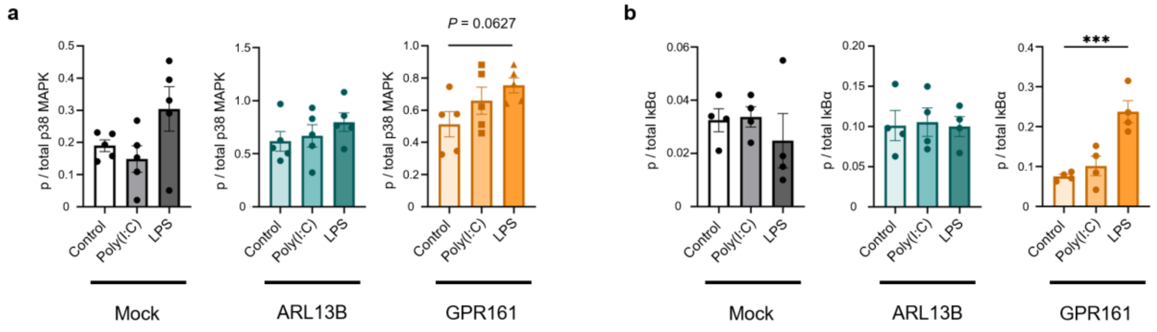


**Supplemental Figure 5. Distinct and shared chromatin accessibility programs induced by ARL13B and GPR161.** (**a**) Quantification of p38 MAPK activation shown in Fig. 5C, expressed as the ratio of p-p38 MAPK to total p38 MAPK (p-p38/total p38). Statistical significance was assessed by one-way ANOVA followed by Dunnett's multiple comparison test versus the control group. (**b**) Quantification of p-IκBα shown in Figure 5c, expressed as the ratio of p-IκBα to total IκBα (p-IκBα/total IκBα). Statistical significance was assessed by one-way ANOVA followed by Dunnett's multiple comparison test versus the control group. ***$P < 0.001$.

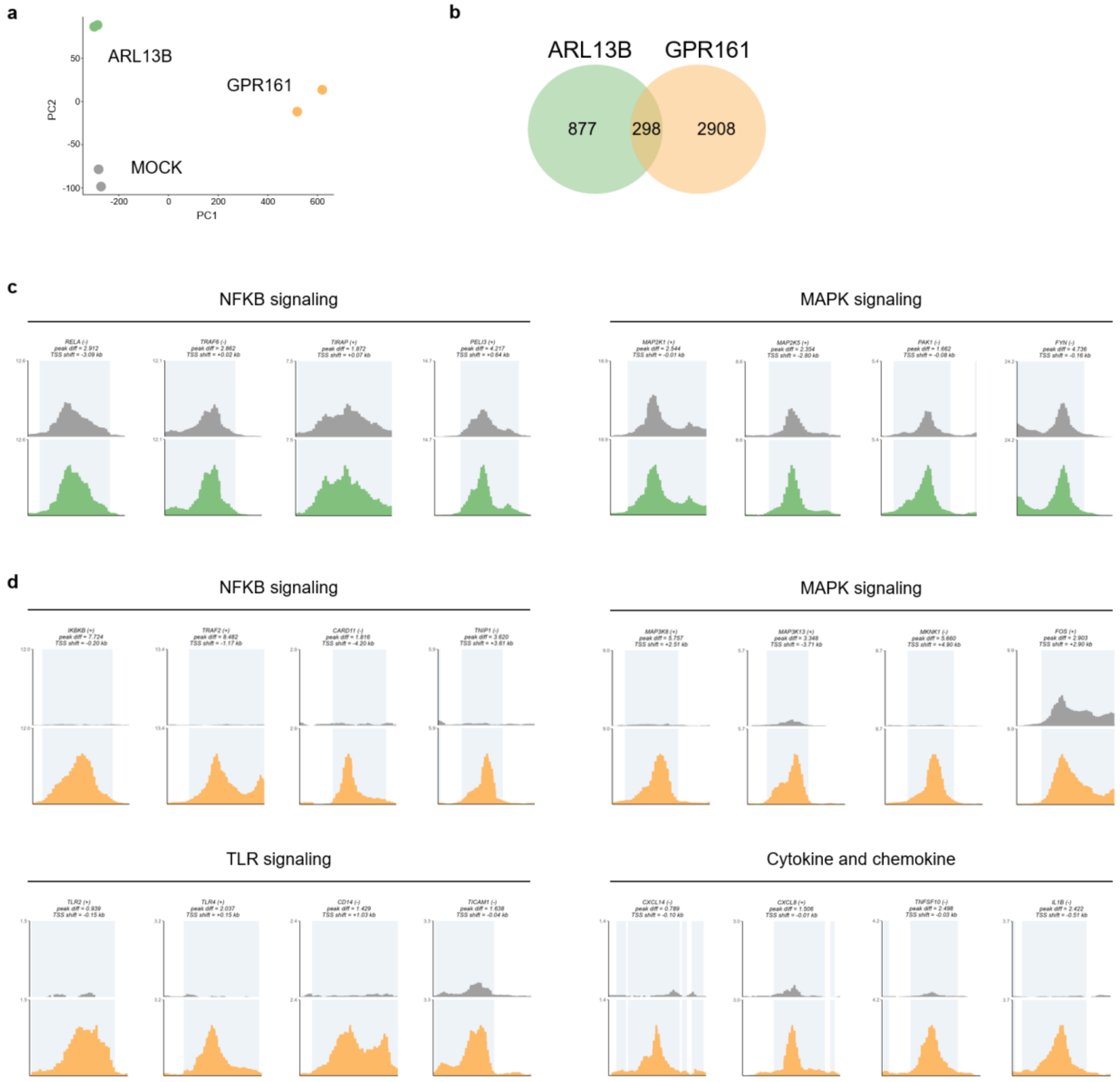


**Supplemental Figure 6. Distinct and shared chromatin accessibility programs induced by ARL13B and GPR161.** (**a**) Principal PCA of ATAC-seq chromatin accessibility profiles across mock-, ARL13B-, and GPR161-overexpressing HaCaT cells. (**b**) Venn diagram showing the overlap of DARs located in the promoter regions of ARL13B- and GPR161-overexpressing HaCaT cells. (**c**) Replicate-pooled ATAC-seq peak tracks showing chromatin accessibility at promoter regions of genes associated with NF-κB (*RELA*, *TRAF6*, *TIRAP*, *PELI3*) and MAPK (*MAP2K1*, *MAP2K5*, *PAK1*, *FYN*) signaling pathways in ARL13B-overexpressing HaCaT cells. Gene orientation is indicated by + (forward strand) and − (reverse strand). (**d**) Replicate-pooled ATAC-seq peak tracks showing chromatin accessibility at promoter regions of genes associated with NF-κB (*IKBKB*, *TRAF2*, *CARD11*, *TNIP1*), MAPK (*MAP3K8*, *MAP3K13*, *MKNK1*, *FOS*), TLR (*TLR2*, *TLR4*, *CD14*, *TICAM1*) signaling pathways, and cytokines/chemokines (*CXCL14*, *CXCL8*, *TNFSF10*, *IL1B*) in GPR161-

overexpressing HaCaT cells. Gene orientation is indicated by + (forward strand) and − (reverse strand).

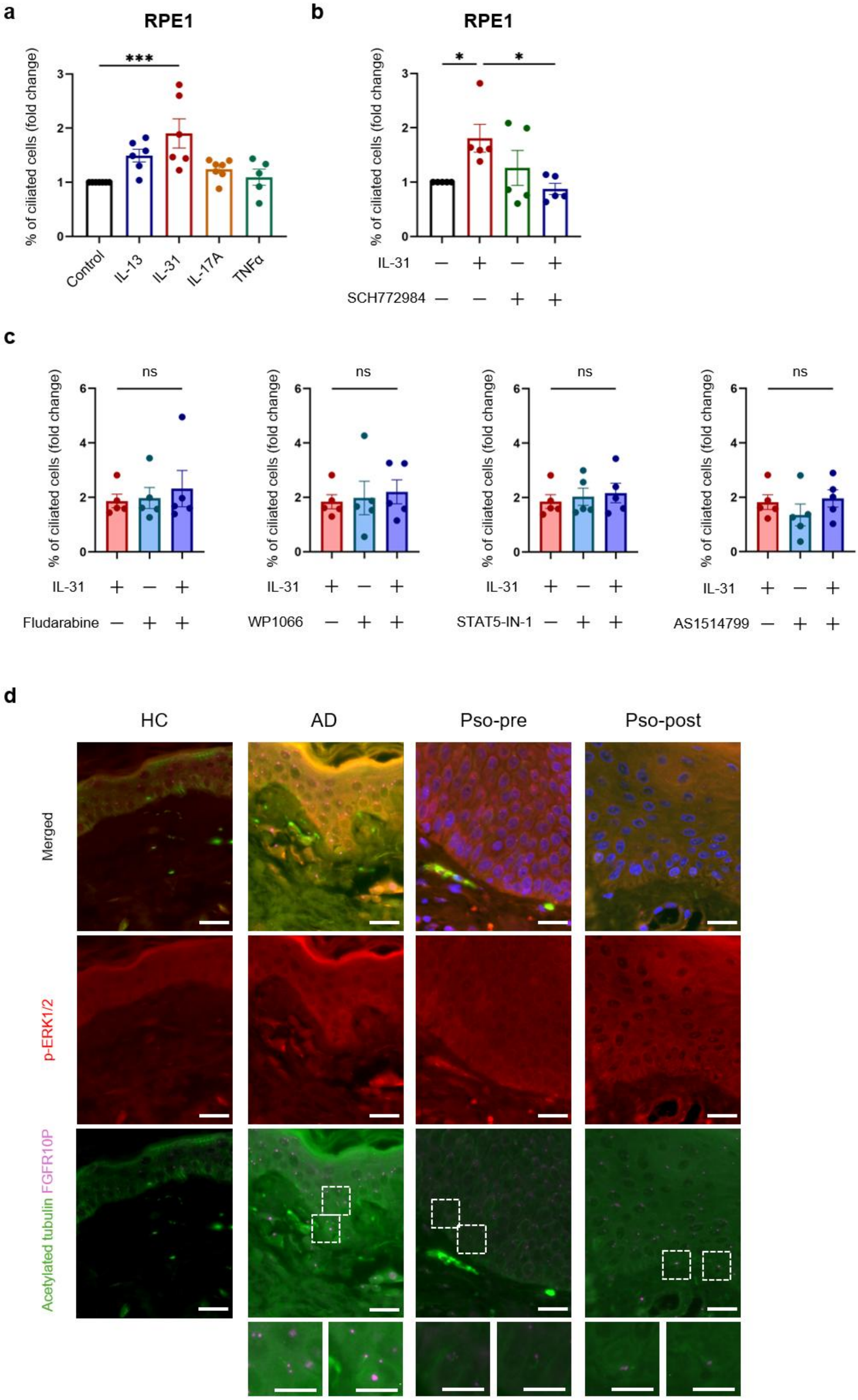
a
RPE1
% of ciliated cells (fold change)
***
Control
IL-13
IL-31
IL-17A
TNFα
b
RPE1
% of ciliated cells (fold change)
*
*
IL-31
SCH772984
c
% of ciliated cells (fold change)
ns
IL-31
Fludarabine
WP1066
STAT5-IN-1
AS1514799
d
HC
AD
Pso-pre
Pso-post
Merged
p-ERK1/2
Acetylated tubulin FGFR1OP

**Supplemental Figure 7. The ERK1/2 pathway promotes IL-31-induced primary cilia formation. (a)** Quantification of ciliated RPE1 cells after cytokine stimulation. Data are presented as fold change relative to the untreated control. **(b)** Quantification of ciliated RPE1 cells after IL-31 stimulation with or without the ERK1/2 specific inhibitor SCH772984. Data are presented as fold change relative to the untreated control. **(c)** Quantification of ciliated RPE1 cells after IL-31 stimulation with or without STAT inhibitors. Statistical significance was assessed using one-way ANOVA, followed by Dunnett's multiple comparison test versus the IL-31 group. **(d)** Representative immunofluorescence images of p-Erk and primary cilia in the skin. Ciliary axonemes were labeled with acetylated α-tubulin (green), centrosomes with FGFR10P (magenta), p-Erk (red), and nuclei were stained with Hoechst 33342 (blue). Scale bar: 20 μm (upper), 10 μm (bottom).
